# Directing cellular transitions on gene graph-enhanced cell state manifold

**DOI:** 10.1101/2024.10.27.620174

**Authors:** Tianze Wang, Yan Pan, Fusong Ju, Shuxin Zheng, Chang Liu, Yaosen Min, Qun Jiang, Xinwei Liu, Huanhuan Xia, Guoqing Liu, Haiguang Liu, Pan Deng

## Abstract

A select few genes act as pivotal drivers in the process of cell state transitions. However, finding key genes involved in different transitions is challenging. To address this problem, we present CellNavi, a deep learning-based framework designed to predict genes that drive cell state transitions. CellNavi builds a driver gene predictor upon a cell state manifold, which captures the intrinsic features of cells by learning from large-scale, high-dimensional transcriptomics data and integrating gene graphs with directional connections. Our analysis shows that CellNavi can accurately predict driver genes for transitions induced by genetic, chemical, and cytokine perturbations across diverse cell types, conditions, and studies. By leveraging a biologically meaningful cell state manifold, it is proficient in tasks involving critical transitions such as cellular differentiation, disease progression, and drug response. CellNavi represents a substantial advancement in driver gene prediction and cell state manipulation, opening new avenues in disease biology and therapeutic discovery.

## Introduction

Understanding the genetic drivers of cellular transitions is crucial for elucidating complex biological processes and disease mechanisms^1–3^. However, identifying these drivers has remained intrinsically challenging due to the sheer number of genes with intricate dependencies involved in transitions, contrasted with our limited experimental capacity and accumulated knowledge. Therefore, in silico methods capable of predicting driver genes across diverse contexts are highly desirable.

Traditionally, efforts to pinpoint critical driver genes have primarily relied on network-based methodologies, with a particular focus on gene regulatory networks (GRNs) ^4–8^. While GRN-centric approaches have made notable progress, they also encounter limitations that hinder their broader use. For example, deducing accurate GRNs within heterogeneous cell populations, which is more relevant to translational research, remains a challenge ^9,10^. Moreover, GRN models tend to prioritize transcription factors and may overlook non-transcriptional drivers of cellular transitions. This limits our understanding of complex cellular processes such as disease progression, immune modulation, and pharmacological responses.

To this end, we develop CellNavi, a deep learning framework designed to predict driver genes and navigate cellular transitions. CellNavi constructs a driver gene predictor on top of a learned manifold that parameterizes valid cell states. This manifold is modeled by mapping raw cell state representations onto a lower-dimensional coordinate space, where the dimensions correspond to intrinsic features of cell states, and the distance reflects the biological similarity between cells. To build this manifold, CellNavi is trained on large-scale, high-dimensional single-cell transcriptomic data, along with prior directional gene graphs that reveal the underlying structure of cell states. By projecting cellular data onto this biologically meaningful space with reduced dimensionality and enhanced biological relevance, CellNavi provides a universal framework that generalizes across diverse cellular contexts, allowing robust driver gene predictions even in previously unexplored cell types or conditions.

Our results show that CellNavi excels at predicting driver genes across a wide range of biological transitions, demonstrating strong performance in quantitative tasks curated in both immortalized cell lines and primary cells. It identifies crucial regulators in T cell differentiation and uncovers key genes associated with neurodegenerative diseases. Notably, CellNavi infers mechanisms of action (MoA) for drug compounds without the need for drug-specific training, underscoring its potential in drug discovery. In summary, CellNavi offers a powerful framework for deciphering cell state transitions and their underlying mechanisms, holding significant promise for advancing cell biology and disease research.

### Overview of CellNavi

CellNavi is designed to predict driver genes for given cellular transitions, where the transcriptomic data of the source and target cells represent the initial and final states of these transitions **(Fig. 1a-c)**.

**Figure 1.**
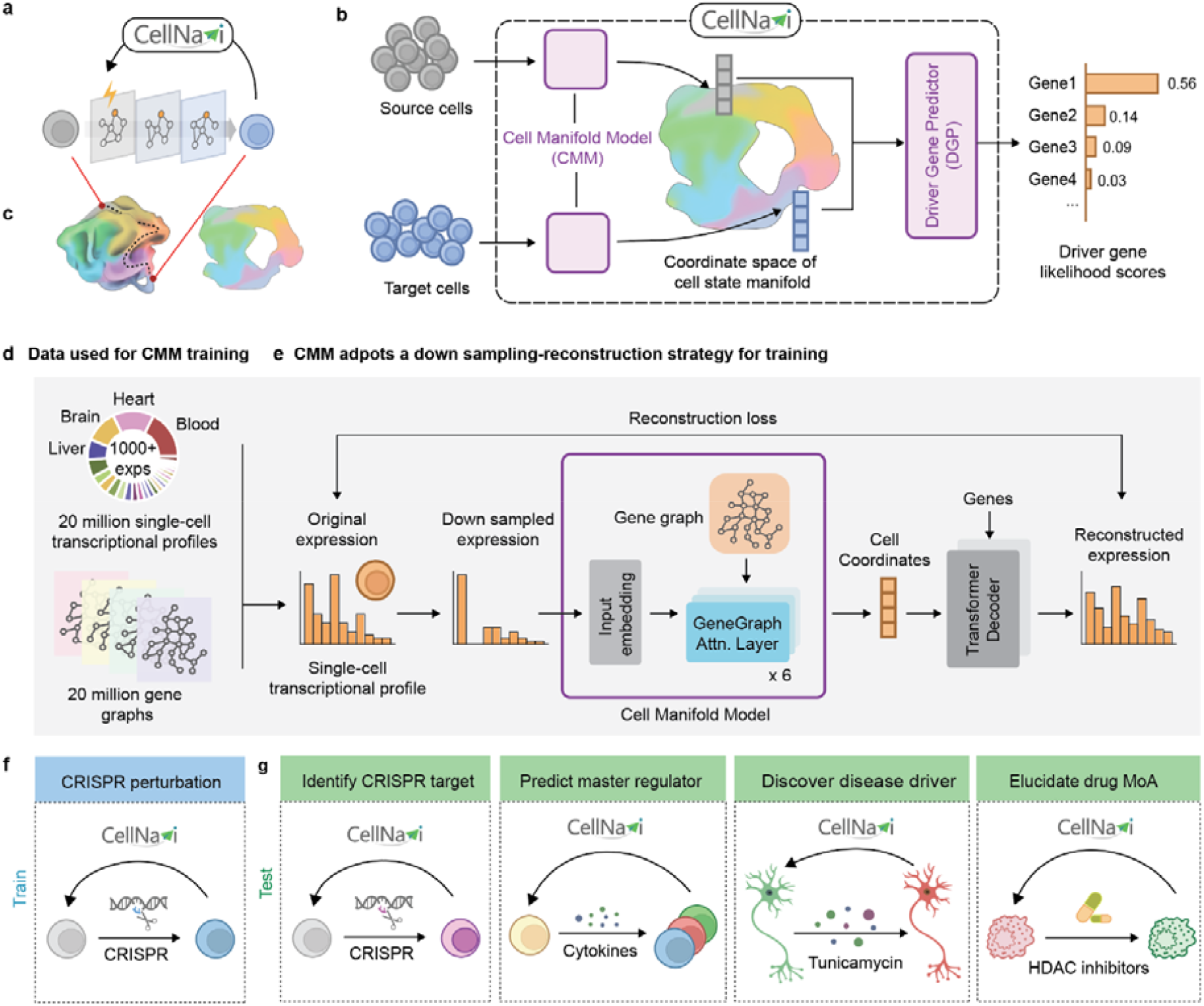
Overview of CellNavi. **a)** Conceptual illustration of CellNavi’s task. Given a pair of source and target cells undergoing a transition induced by stimuli, CellNavi predicts the driver gene responsible for this transition. **b)** Workflow of CellNavi. The Cell Manifold Model (CMM) maps the source and target cells onto a coordinate space of the cell manifold. The Driver Gene Predictor (DGP) then uses the cell coordinates produced by the CMM to rank the candidate genes by likelihood scores. **c)** Illustration of the cell manifold and its coordinate space. **d)** Data used for the CMM training. **e)** Training of the CMM. The CMM consists of six GeneGraph Attention (Attn.) layers designed to incorporate graph-based information. During training, single-cell transcriptomic profiles are randomly sampled from the curated HCA dataset and used as input. Cell embeddings generated by the model are then used by a Transformer decoder to reconstruct gene expression profiles. **f)** Data used for the DGP training. **g)** Application scenarios and test cases of CellNavi. Schematic elements created with BioRender.com.

CellNavi comprises two main components: the Cell Manifold Model (CMM), which captures and represents cell states, and the Driver Gene Predictor (DGP), which identifies key genes driving these transitions based on learned cell representations **(Fig. 1b)**.

The CMM is built to capture valid cell states across diverse biological contexts. While transcriptomes are often used to represent cell states, valid cell states do not span the entire high-dimensional transcriptomic space but instead form a lower-dimensional manifold **(Fig. 1c)**. To model this, the CMM maps transcriptomic vectors to a lower-dimensional coordinate space that represents the intrinsic features of cell states, while preserving the relative similarities between cells (dimensionality considerations are discussed in **Supplementary Note 1**).

We first curated a dataset of approximately 20 million single-cell transcriptomic profiles sourced from the Human Cell Atlas^11^ **(Fig. 1d)** and adapted a Transformer architecture based on attention mechanisms, known for its ability to discern complex patterns in large-scale data^12–17^, to train the CMM **(Fig. 1e)**. The training involved a self-supervised down-sampling-reconstruction task **(Methods and Supplementary Note 2)**. To prioritize cell rather than gene-level representations, we developed a decoder module to reconstruct gene expression profiles from the cell coordinates—representations of cells within the coordinate space of the cell state manifold—generated by the CMM **(Fig. 1e and Extended Data Fig.1, Methods)**. This approach aligns cells across varying sequencing depths **(Extended Data Fig. 2)** and recapitulates developmental trajectories from single cells **(Extended Data Fig. 3)**, indicating that it captures both intra- and inter-cellular features.

However, relying solely on transcriptomic data may overlook the intricate gene-gene interactions that are crucial for describing and distinguishing cell states. To address this, we incorporated 20 million cell-specific gene graphs into the CMM training process **(Figs. 1d-e)**. These graphs encode directional connections derived from a prior network that spans over 30,000 human genes and their associated signaling pathways^18^ **(Methods)**. To be more specific, in these gene graphs, each edge represents a causal relationship between two genes, with the direction of the edge indicating the regulatory influence **(Methods)**. These graphs provide richer information about the complex dependencies among genes, which extend beyond simple transcriptomic data, hence better implying intrinsic variables spanning the valid cell space. To leverage these gene graphs, we replaced the standard Transformer encoder layer in the CMM using a GeneGraph Attention Layer **(Extended Data Fig. 1b)**. These layers, inspired by attention variants tailored for graph data^19^, can process gene networks, thus enabling the model to integrate critical gene-gene relationships. With these designs, the model is driven to cultivate a manifold that systematically represents cell states and effectively reflects the relationships between cells, forming an informative foundation for driver gene prediction.

Building upon this manifold, we developed the DGP to predict genes driving specified cellular transitions **(Extended Data Fig. 4, Methods)**. The DGP is trained on CRISPR screen data, which link genetic perturbations to consequent changes in cell states^20–25^. We designated unperturbed controls and CRISPR-perturbed cells as source and target pairs, respectively, and utilized validated perturbed genes as labels for joint training (fine-tuning) of the CMM and DGP **(Fig. 1f)**. Specifically, for each cell pair, their transcriptomic profiles are transformed into cell coordinates by the CMM, which are then processed by the DGP to generate a likelihood score vector indicating the probability that various candidate genes are orchestrating the transitions **(Extended Data Fig. 4)**.

We demonstrate that CellNavi, fine-tuned on CRISPR screen data—typically conducted on cultured cells or homogeneous populations and focusing on immediate genetic perturbations—can be extended to more complex transitions in heterogeneous tissues and primary cells (**Fig. 1g and Extended Data Fig. 5**). By leveraging a biologically meaningful manifold, CellNavi generalizes knowledge gained from CRISPR screens beyond their original scope, to cellular transitions that are challenging to investigate using regular CRISPR methodologies. However, we acknowledge that CellNavi’s performance in specific contexts may benefit from additional fine-tuning on relevant CRISPR datasets. Incorporating expanded experimental data may further enhance its applicability across diverse biological settings with minimal adaptation.

### Quantitative evaluation of CellNavi

To assess the capabilities of CellNavi, we first evaluated its performance on CRISPR perturbation datasets, where driver gene information is well-established for transitions from source (unperturbed) to target (perturbed) cells.

We initially applied CellNavi to the Schmidt dataset, a CRISPR activation screen profiling 69 genetic perturbations^26^. This dataset captures distinct expression profiles and molecular phenotypes across both resting and re-stimulated T cells, within and between different cell types, before and after perturbations **(Supplementary Fig. 1)**. We fine-tuned our model on re-stimulated T cells and tested it on resting T cells **(Fig. 2a)**. This setup allowed us to evaluate CellNavi’s ability to generalize across heterogeneous primary cells and predict driver genes in new cell states.

**Figure 2.**
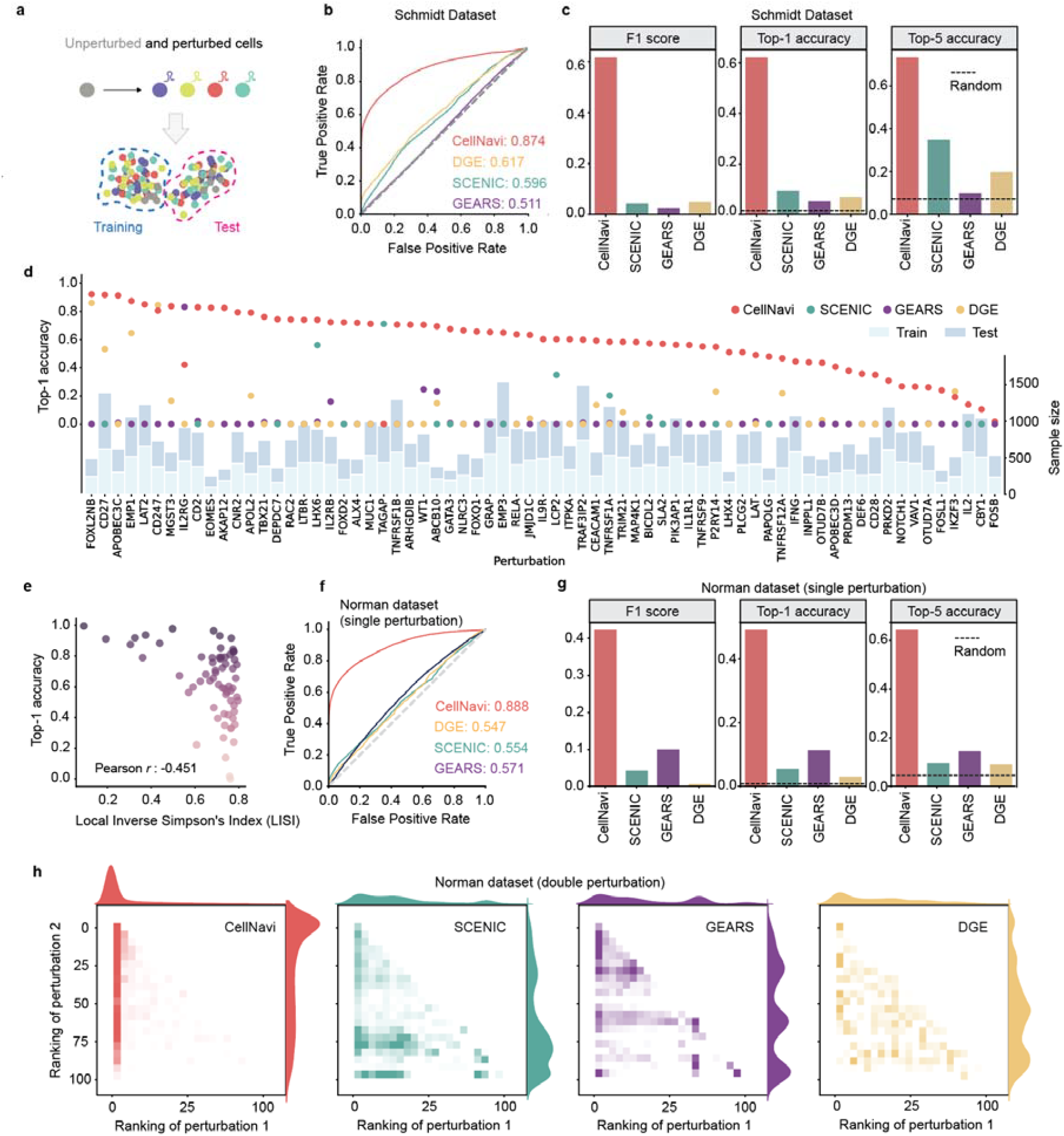
Quantitative assessment of CellNavi. **a)** Schematic of the quantitative evaluation framework. CRISPR-perturbed cells and their unperturbed controls are used for model training and evaluation, with data split by cell states to enable more rigorous testing. **b)** top-1 accuracy, top-5 accuracy, and F1 score for driver gene prediction in the Schmidt dataset, comparing CellNavi with alternative methods. The dashed line indicates the performance of a random guess. **c)** AUROC scores for driver gene prediction in the Schmidt dataset, comparing CellNavi with alternative methods. **d)** Average top-1 accuracy for each gene. Left y-axis: top-1 accuracy of different methods for each gene. Right y-axis: the number of training (light blue) and test (steel blue) samples. **e)** Negative correlation between CellNavi’s top-1 accuracy and the average LISI score across genes (Pearson correlation coefficient: -0.451). A LISI score of 1 indicates indistinguishable perturbation effects, while a score of 0 suggests a distinct perturbation pattern. Dot colors represent the top-1 accuracy for individual genes. **f)** top-1 accuracy, top-5 accuracy, and F1 score for driver gene prediction in the Norman dataset (single perturbation), comparing CellNavi with alternative methods. The dashed line indicates the performance of a random guess. **g)** AUROC scores for driver gene prediction in the Norman dataset (single perturbation), comparing CellNavi with alternative methods. **h)** Distribution of predicted rankings for perturbed gene pairs. “Perturbation 1” represents genes ranked higher, and “Perturbation 2” represents genes ranked lower. *n* = 4,916. Source data for panels b, c, f, g are available in Supplementary Table 1.

For each source-target cell pair, CellNavi prioritizes candidate genes based on their predicted likelihood scores. Across 23,047 source-target cell pairs, CellNavi achieves a top-1 accuracy of 0.621 and a top-5 accuracy of 0.733 **(Fig. 2b)**, while maintaining strong performance across additional metrics **(Figs. 2b-c and Extended Data Fig. 6)**. Interestingly, substantial variation in top-1 accuracy was observed across perturbed genes, independent of sample size **(Fig. 2d)**. Correlation analysis between gene-wise performance and the Local Inverse Simpson’s Index (LISI) ^27^ suggests that CellNavi’s accuracy is influenced by the degree of perturbation heterogeneity: perturbations with low average LISI values, indicative of a more distinct and homogeneous response, were associated with higher accuracy (top-1 accuracy > 0.8, **Fig. 2e**).

To demonstrate CellNavi’s effectiveness, we compared it to two alternative methods: SCENIC/SCENIC+^4,5^, a training-free approach that infers GRNs from transcriptomic data with a focus on master regulators, and GEARS^28^, an in silico perturbation approach, which targets a partially inverse problem of cellular transition prediction **(Methods)**. Both SCENIC and GEARS exhibited significantly lower performance compared to CellNavi **(Figs. 2b-d and Extended Data Fig. 6a)**. In addition, SCENIC, the network-based approaches, faced challenges in identifying regulons at the single-cell level **(Supplementary Fig. 2)**, and therefore struggled to make predictions in many cases. To investigate whether this is a broad challenge for GRN inference methods, we evaluated three alternative GRN inference approaches: GENIE3^29^, GRNBoost2^30^, and RENGE^31^ **(Methods)**. These methods similarly exhibited poor performance in single-cell contexts **(Supplementary Table 1)**.

CellNavi does not simply predict driver genes from expression changes. We conducted an ablation study by systematically removing the expression of perturbed genes from the input. Although this led to a decrease in performance, CellNavi still maintained substantial predictive accuracy, far surpassing expectations of random prediction **(Extended Data Fig. 7)**. In addition, differential gene expression (DGE) analysis revealed that the rankings of differentially expressed genes were poorly correlated with the actual perturbed genes **(Figs. 2b-d and Extended Data Fig. 6a)**. These results suggest that CellNavi identifies driver genes beyond those detectable by expression shifts alone.

We further tested CellNavi on the Norman dataset^32^, which features a CRISPR interference screen on the K562 cell line. This dataset encompasses 105 single-gene and 131 gene pair perturbations, allowing us to assess CellNavi’s performance on transitions driven by both single and multiple genes. Using the unsupervised Leiden algorithm^33^, we stratified the cells by cluster, holding out one cluster for testing and training on the remaining ones **(Supplementary Fig. 3 and Fig. 2a)**. To ensure rigorous evaluation, we excluded all multi-gene perturbations from training.

CellNavi maintained strong performance on single driver gene prediction in the Norman dataset **(Figs. 2f-g, Extended Data Fig. 6b and Supplementary Table 1)**. To evaluate multi-gene scenarios, we focused on the predicted rankings of perturbed genes. CellNavi ranked the first and second perturbed genes at averages of 7.9 and 31.2 out of 105 candidates, respectively, significantly outperforming all other tested methods **(Fig. 2h)**.

Several recent studies have indicated that linear models can outperform deep learning methods in cell modeling tasks^34–37^. To investigate this, we evaluated multiple linear models for driver gene prediction under various conditions. Our results showed that CellNavi consistently outperformed these linear models by a substantial margin across settings (**Supplementary Note 3**). Furthermore, we applied cross-validation to ensure robust and unbiased evaluation and found that CellNavi demonstrated consistently superior performance across these conditions **(Supplementary Tables 2-3)**. Altogether, these results, spanning diverse datasets and metrics, highlight CellNavi’s strong capability to identify genes driving cellular changes, even in previously uncharacterized cell states.

### Evaluating model components and graph configurations

To assess the contributions of the CMM and DGP components, and to evaluate whether pretraining with the CMM improves generalization across biological contexts, we designed two ablated methods. The first combined the DGP with raw gene expression vectors instead of outputs from the CMM (no-CMM). The second replaced the DGP with a simpler multinomial logistic regression model (no-DGP). In addition to the Norman single perturbation split, which utilizes a cluster-based holdout strategy (out-of-domain split), we curated an alternative evaluation approach using random holdout to simulate a scenario without generalization (in-domain split). Removing either CMM pretraining (no-CMM) or DGP fine-tuning (no-DGP) led to reduced performance; however, for out-of-domain split, the absence of CMM pretraining (no-CMM) caused a greater drop in performance compared to the in-domain split scenario (**Extended Data Fig. 8**). These results highlight that CMM pretraining is essential for generalization across biologically diverse contexts, while DGP fine-tuning further optimizes task-specific predictions.

We also evaluated the impact of the NicheNet gene graph on CellNavi’s predictions. Replacing NicheNet with GRNs inferred using GENIE3, GRNBoost2, or RENGE resulted in reduced performance (**Extended Data Table 1**), underscoring the advantage of integrating pathway-level information beyond GRNs, particularly in modeling perturbation-induced transitions. Furthermore, we tested graph configurations with varying levels of connectivity, including fully connected graphs, sparsified graphs with edges reduced to 1/10 or 1/20 of the original graph, and random graphs with the same sparsity as NicheNet (**Methods**). All alternative configurations led to further performance declines relative to biologically meaningful graphs constructed using diverse GRN inference methods (**Extended Data Table 1**). Collectively, these results emphasize the importance of leveraging biologically meaningful and comprehensive gene graphs, such as NicheNet, to ensure predictive robustness and accuracy.

### CellNavi identifies key genes in T cell differentiation

We next applied CellNavi to the Cano-Gomez dataset^38^, which profiled T cell differentiation by stimulating naïve and memory CD4+ T cells in vitro with anti-CD3/anti-CD28 and cytokines. During this process, external signals, such as antigens and cytokines, activate key genes modulating genetic circuits and gene expression programs, allowing T cells to adopt specialized functions. We assessed whether CellNavi could identify such key genes underlying transitions.

For this dataset, we constructed source-target cell pairs using Th0 cells as the source and cytokine-induced cells as targets. As cells differentiated into various effector T cell subtypes after stimulation^26,38–43^, we first compiled a comprehensive marker gene set and computed a “transition score” to quantify differentiation into these subtypes for each cell. Notably, marker genes associated with IL2-high, IFNγ-high, and Th2 cells were strongly enriched **(Supplementary Fig. 4)**, and transition scores towards these cell types demonstrated clear patterns **(Figs. 3a–c, Methods)**. We then examined CellNavi’s ability to identify driver genes across these effector T cell groups. Corresponding cell pairs were input into a CellNavi model trained on the Schmidt dataset, which encompasses extensive immune-related gene programs. Finally, we curated a literature-based list of established driver genes for phenotypic transitions towards specific effector cell types^26,42–49^ **(Supplementary Table 4)**, and evaluated CellNavi’s performance in prioritizing these genes.

**Figure 3.**
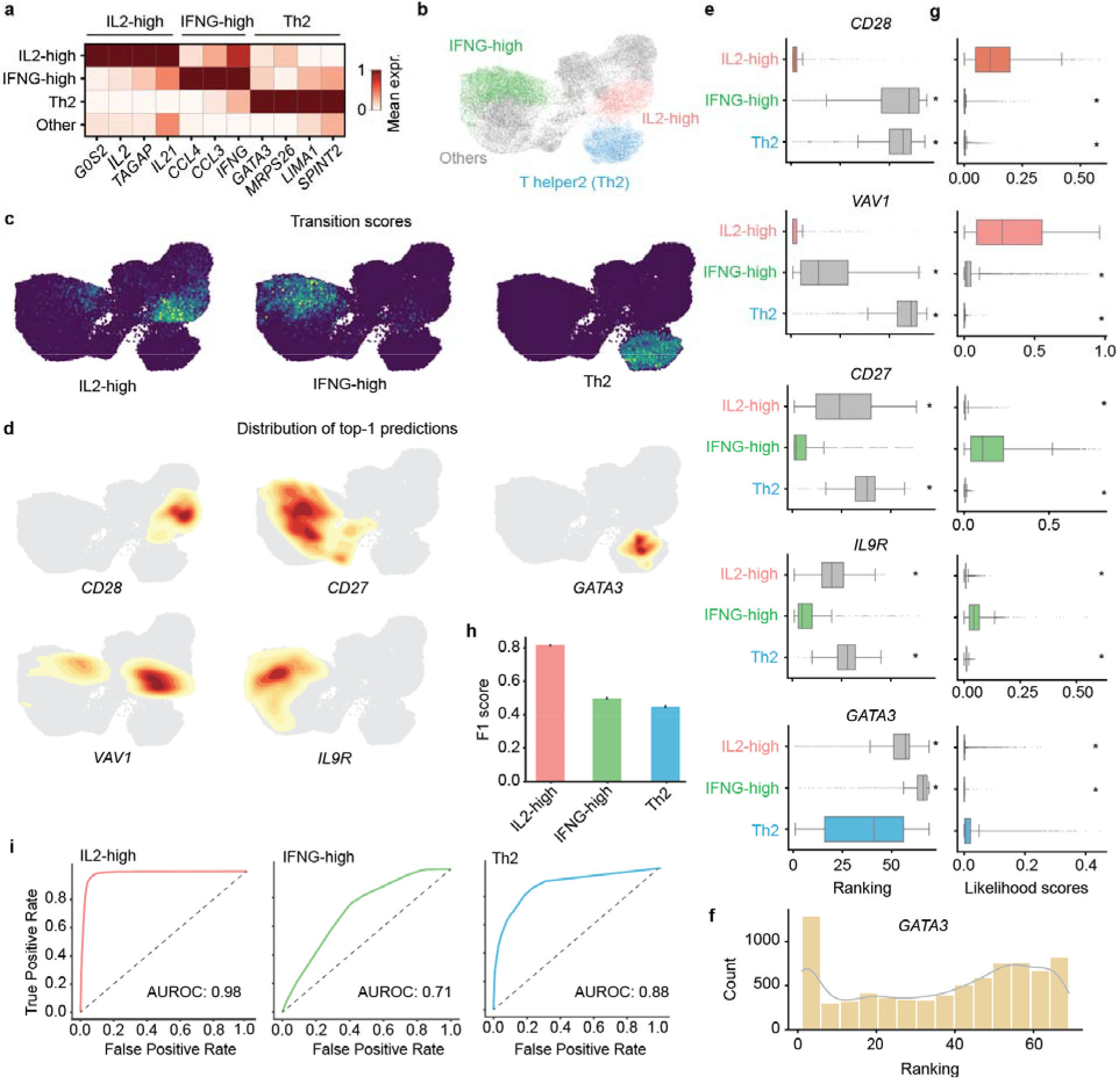
CellNavi identifies key genes involved in T cell differentiation. **a)** Changes in expression levels of canonical marker genes corresponding to specific T cell groups. **b)** UMAP visualization of source-target T cell pairs, colored by effector T cell groups classified based on transition scores. Each data point represents a source-target cell pair representation generated by CellNavi. **c)** Transition scores calculated using IL2-high, IFNγ-high, and Th2-related marker genes referenced in panel a. **d)** Distributions of established driver genes predicted by CellNavi for IL2-high cells (*CD28, VAV1*), IFNγ-high cells (*CD27, IL9R*), and Th2 cells (*GATA3*). **e)** Predicted rankings of established driver genes across different cell groups. Center line, median; box limits, upper and lower quartiles; whiskers, 1.5x interquartile range; points, outliers. *n* = 23,342. *P*-values were calculated with two-sided Mann–Whitney *U* test. Star (*) indicates a *p*-value smaller than 1e-6. **f)** Distribution of predicted rankings for *GATA3* in Th2 cells. **g)** Predicted likelihood scores for established driver genes in different cell groups. Center line, median; box limits, upper and lower quartiles; whiskers, 1.5x interquartile range; points, outliers. *n* = 23,342. *P*-values were calculated with two-sided Mann–Whitney *U* test. Star (*) indicates a *p*-value smaller than 1e-6. **h)** F1 scores for predicting effector T cell types using likelihood scores. Center: mean. Error bar: standard error, calculated from ten-fold cross-validation (Methods). *n* = 10. **i)** AUROC scores for predicting effector T cell types using likelihood scores (Methods).

CellNavi accurately ranked *CD28* and *VAV1*, key drivers of IL2-high cells, as the top candidates in the IL2-high group defined by the transition score **(Fig. 3d)**. Similarly, high rankings were observed for *CD27* and *IL9R* in IFNγ-high cells, and *GATA3* in Th2 cells **(Fig. 3d)**. We further analyzed the average rankings of these established driver genes across the different effector cell groups. As expected, the relevant driver genes consistently ranked higher in their corresponding cell groups where they are known to drive differentiation. Notably, *CD28, VAV1, CD27*, and *IL9R* achieved average rankings of 2.6, 2.9, 5.3, and 8.5, respectively, in their associated cell groups, significantly outperforming their rankings in unrelated groups **(Fig. 3e)**. These results demonstrate CellNavi’s effectiveness in identifying key genes that govern distinct differentiation pathways while distinguishing between cell fates. However, *CD28*’s dual role in IL2 and IFNγ regulation was not fully captured by the model. Additionally, although *GATA3* ranked highly in Th2 cells, its average ranking was not as strong as expected. Upon further inspection of the Th2 cluster, we observed that *GATA3* was ranked first in an aggregated subset of cells, while its ranking was more dispersed across the entire Th2 group **(Fig. 3f)**, suggesting heterogeneity within the cluster.

Next, we examined the likelihood scores assigned by CellNavi to driver genes across different cell groups. For known driver genes, CellNavi consistently assigned higher likelihood scores within their corresponding cell groups compared to other groups **(Fig. 3g)**, suggesting that these scores accurately prioritize key driver genes. Additionally, the scores could be used to distinguish cell states undergoing specific transitions **(Fig. 3h-i, Methods)**, offering an alternative approach for cell state characterization.

### CellNavi predicts key genes during pathogenesis

We then investigated whether CellNavi could predict key genes involved in disease progression, using an *in vitro* model system of neurodegenerative diseases, specifically the Fernandes dataset^50^. This system comprises induced pluripotent stem cells (iPSC)-derived dopaminergic neurons subjected to Tunicamycin treatment. Tunicamycin induces endoplasmic reticulum (ER) stress and Parkinson’s disease (PD)-like symptoms by inhibiting N-linked glycosylation^51^, a process that affects a broad spectrum of proteins post-translationally, without perturbing any single gene directly.

Prior to this analysis, CellNavi was trained on single-cell CRISPR screen data on iPSC-derived neurons from a different study, the Tian dataset^52^. While both studies investigate neurodegenerative diseases using human iPSC-derived neurons, they differ in the source of iPSCs and the differentiation protocols, resulting in the generation of distinct neuron types^50,52,53^ **(Fig. 4a)**.

**Figure 4.**
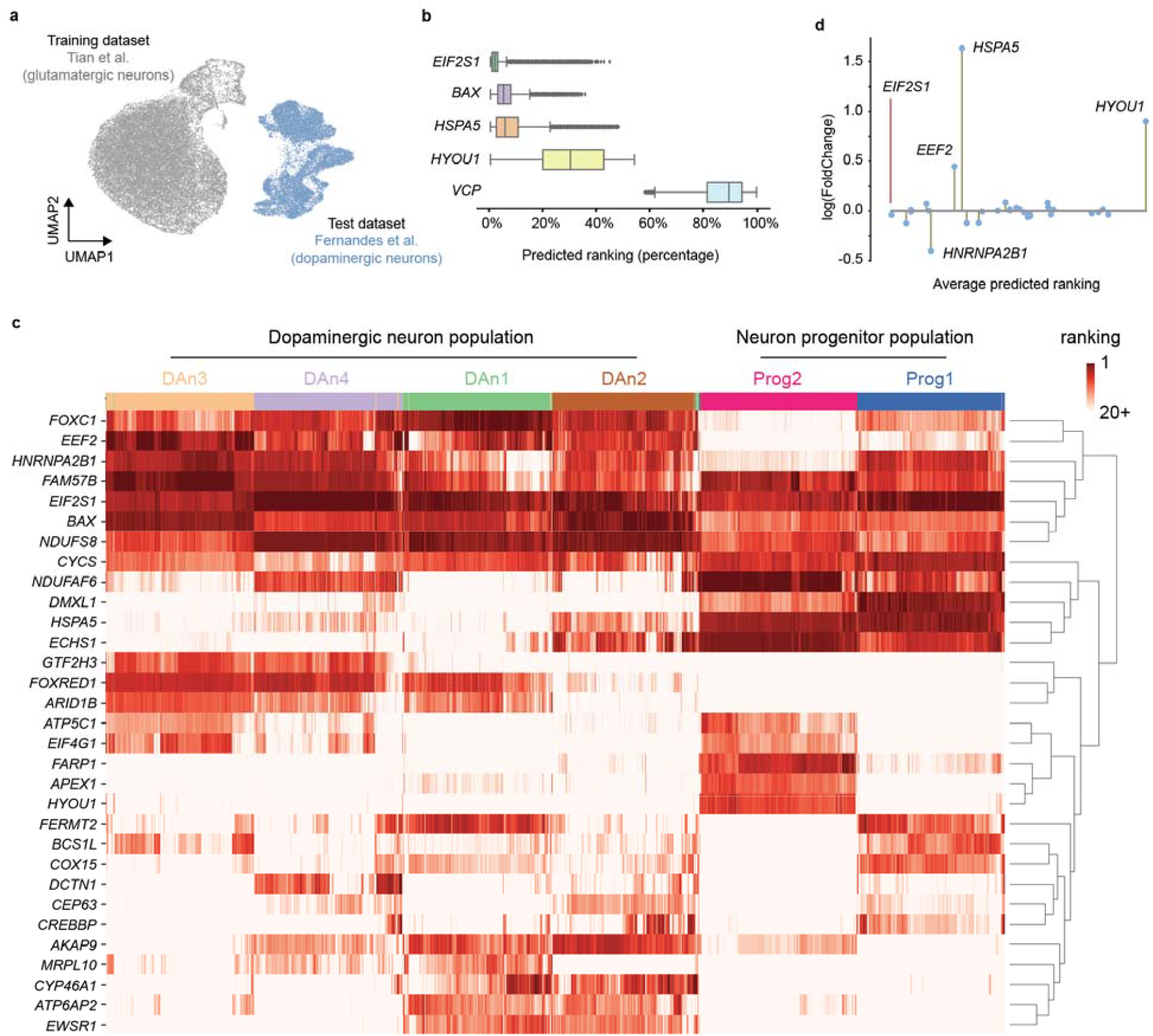
CellNavi predicts key genes involved in neurodegenerative pathogenesis. **a)** UMAP visualization of transcriptomic profiles from neurodegenerative disease-related datasets. Gray: iPSC-derived glutamatergic neurons (Tian dataset) used for model training. Blue: iPSC-derived dopaminergi neurons (Fernandes dataset). **b)** Predicted rankings for ER stress response-associated genes, based on likelihood score vectors generated by CellNavi. Center line, median; box limits, upper and lower quartiles; whiskers, 1.5x interquartile range; points, outliers. *n* = 47,437 **c)** Distribution of the top 20 predicted gene across all cell pairs. Rows represent cell types as defined by the original publication (Fernandes et al.). Darker colors indicate higher rankings, and lighter colors indicate lower rankings. Hierarchical clustering was performed using Ward’s method. **d)** Expression changes for the top 20 predicted genes. The x-axi shows the average ranking of each gene across cell pairs, while the y-axis indicates the fold change in expression between target cells and source cells.

After training, we input approximately 47,000 source-target cell pairs from the Fernandes dataset into CellNavi, using untreated cells as sources and cells exposed to tunicamycin as targets. We asked CellNavi to prioritize 184 candidate genes, including 5 known ER stress response genes. CellNavi successfully pinpointed *EIF2S1, BAX*, and *HSPA5*, which achieved median rankings of 3, 7, and 16, respectively, among the candidate genes **(Fig. 4b)**. However, *HYOU1* and *VCP* ranked lower. One possible explanation is that these genes play more nuanced roles in the ER stress response or are involved in pathways not prominently activated under the specific experimental conditions of this study.

We next examined the top 20 predicted genes for each cell pair. While a total of 31 genes were significantly enriched **(Fig. 4c)**, *FAM57B, EIF2S1, NDUFS8, BAX*, and *CYCS* consistently ranked highest across the majority of cells. Notably, *EIF2S1* and *BAX* are well-established ER stress regulators, while *NDUFS8* and *CYCS* are linked to mitochondrial stress, which is often closely associated with ER stress^54^. In parallel, Fernandes et al. previously identified six subtypes of iPSC-derived neurons from transcriptomic data and our top 20 predictions revealed subtype-specific gene preferences. For instance, our model suggests that *FARP1, CELF1, HYOU1*, and *APEX1* may play more critical roles in progenitor cells **(Fig. 4c)**. Lastly, except for *HSPA5* and *HYOU1*, most predicted genes showed modest expression changes **(Fig. 4d and Supplementary Fig. 5)**, consistent with previous observations that CellNavi identifies key regulators beyond those detectable by expression shifts alone.

### CellNavi reveals mechanisms of action for drug compounds

Understanding the mechanisms of action (MoA) of novel drug candidates may enhance drug safety and efficacy, reduce development costs, and accelerate drug discovery process. However, conventional drug screening paradigms often fall short in elucidating the cellular-level effects that drive biological functions and therapeutic outcomes.

Here, we applied CellNavi to predict key genes modulated by Histone Deacetylase (HDAC) inhibitors, a class of anti-tumor drugs with promising therapeutic potential in cancer treatment^55^. HDACs are enzymes integral to post-translational protein modifications and interact with various oncogenic pathways to promote tumor progression^56,57^. The intricate downstream pathways influenced by HDAC presents a significant challenge in fully understanding mechanisms through which HDAC inhibitors exert their effects within cells.

For this purpose, we applied CellNavi to a chemical screen that quantified the transcriptomic response of K562 cells to 17 distinct HDAC inhibitors (referred to as the Srivastan dataset)^58^. In this setup, vehicle-treated cells were designated as sources, while cells exposed to the HDAC inhibitors served as targets. The predicted likelihood score indicated whether a gene was modulated during drug treatment, with higher scores suggesting a more prominent role during treatment with specific HDAC inhibitors. To be noted, CellNavi was trained exclusively on genetic perturbations^25^.

While the transcriptomic data depicted a mixed response across the inhibitors **(Supplementary Fig. 6)**, the likelihood score vectors effectively clustered the inhibitors into distinct clusters **(Figs. 5a-b and Supplementary Fig. 7)**. Further analysis revealed diversity in the top-ranked driver genes **(Fig. 5c)**. Specifically, cells treated with mocetinostat, tucidinostat, entinostat, and tacedinaline (grouped in cluster 3) exhibited high scores for mitochondrial-related genes such as *MRPS31* and *NDUFB7*. In contrast, most other compounds prioritized genes related to RNA splicing and transcription regulation, such as *PRPF3* and *POLR2A*.

**Figure 5.**
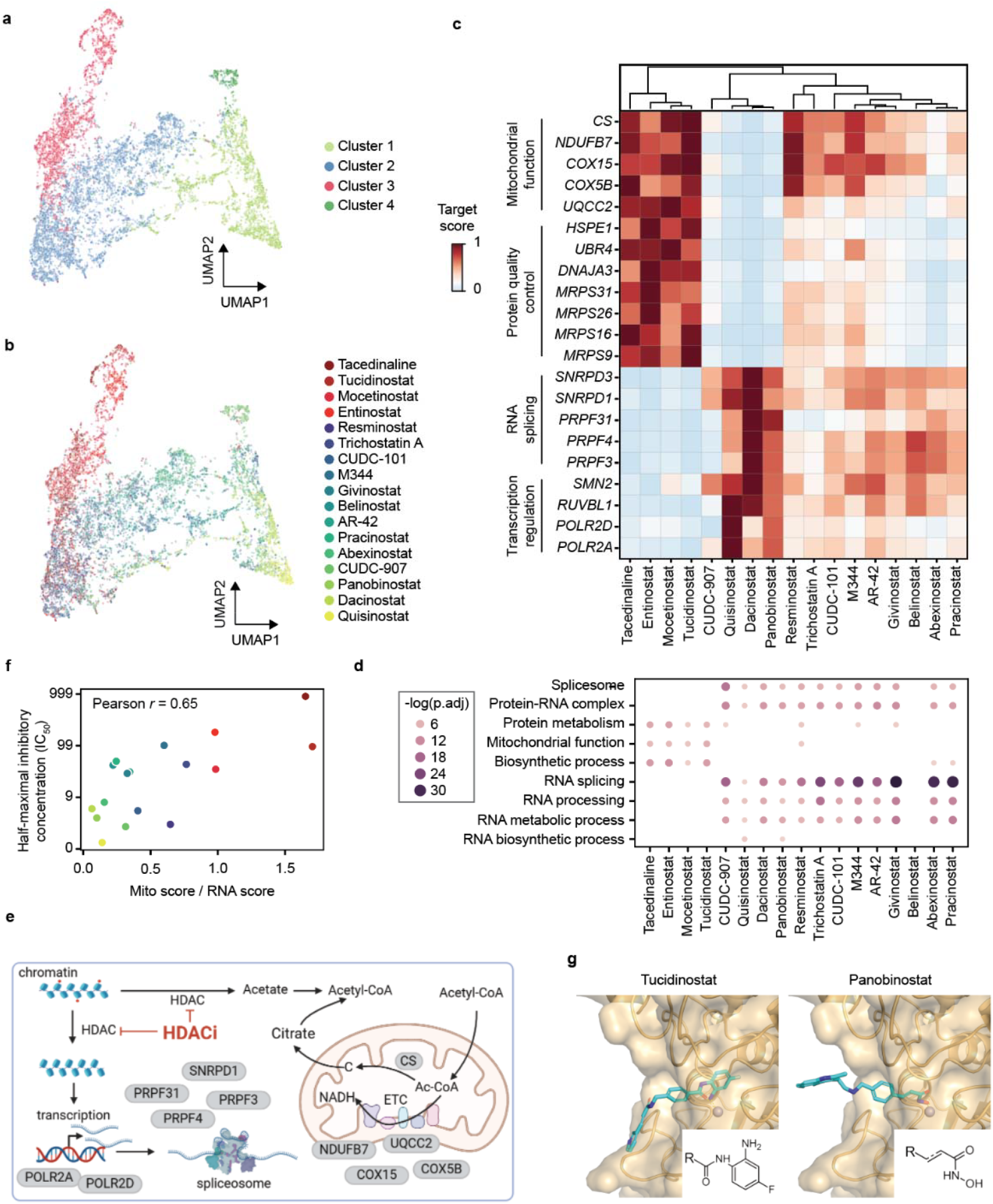
CellNavi reveals diverse downstream gene programs affected by HDAC inhibitors. **a-b)** UMAP visualization of cells treated with HDAC inhibitors. Each cell is represented as a 2,057-dimensional vector consisting of likelihood scores predicted by CellNavi for each candidate driver gene. In a, cells are colored by clusters identified using the Leiden algorithm, and in b, by HDAC inhibitor type. **c)** Average likelihood scores for top-ranked genes in each treatment group, with hierarchical clustering performed using Ward’s method. **d)** Gene Ontology (GO) enrichment analysis for each treatment group. The size and darkness of the dots correlate negatively with the adjusted p-value (one-sided Fisher’s exact test with Benjamini–Hochberg correction for multiple comparisons). See Supplementary Fig. 8 for a complete list of GO enrichment results. **e)** Schematic representation of HDAC inhibitor mechanisms. Proteins encoded by top-ranked driver genes are shown in gray, and red dots on chromatin indicate histone acetylation. ETC: electron transport chain. Ac-CoA: acetyl-CoA. Diagram created with BioRender.com. **f)** Scatter plot showing the correlation between IC50 values and the functional selectivity predicted by CellNavi. Each dot represents a compound, colored according to the clusters in b. Mito score: averaged likelihood scores for genes involved in mitochondrial functions. RNA score: averaged likelihood scores for genes involved in RNA regulation. **g)** Binding modes of tucidinostat and panobinostat at the active site of the zinc-dependent HDAC2 enzyme, with the enzyme represented as a surface representation and the drug compounds in stick representation. Shared warhead (or structural) motifs of different compound classes are highlighted in the bottom right corner. See Supplementary Fig. 8 for a complete list of molecular structures.

Gene Ontology (GO) enrichment analysis of the top 50 genes predicted for each inhibitor revealed a consistent pattern **(Fig. 5d and Supplementary Fig. 8)**: compounds in cluster 3 were enriched for genes involved in biosynthetic processes, mitochondrial function, and protein metabolism, whereas compounds in other clusters were enriched in gene programs related to RNA splicing, processing, and metabolism. These findings align with the known effect of deacetylation inhibition, which lowers cytoplasmic acetate levels and alters acetyl-CoA concentrations, a key metabolite involved in cellular metabolism^58^. Moreover, the results suggest that certain HDAC inhibitors may preferentially target chromatin regions regulating RNA processing genes, which are crucial for the tumor cell proliferation ^59–61^ **(Fig. 5e)**.

Intriguingly, we observed a correlation between the selectivity of downstream gene programs and the half-maximal inhibitory concentration (IC_50_) values reported in the literature^58^ **(Fig. 5f)**. Specifically, compounds with lower IC_50_ values tend to influence RNA-related pathways, whereas those with higher IC_50_ values were associated with mitochondrial functions. To further explore the molecular basis of this divergence, we examined the interactions between human HDAC2 and either panobinostat (enriched for RNA-related genes) or tucidinostat (enriched for mitochondrial-related genes). Although molecular docking revealed no major differences in their potential interactions with the zinc-dependent HDAC protein, the aniline group in tucidinostat allowed it to embed more deeply into the HDAC2 pocket (**Fig. 5g**). Interestingly, all four compounds in cluster 3 shared similar warheads, a feature absent in other compounds **(Fig. 5g and Supplementary Fig. 9)**. This structural feature introduces a steric effect that may influence the efficacy of compounds^62^ and lead to divergent downstream response, a phenomenon known as functional selectivity^63–66^. However, the mitochondrial preference and lower potency of compounds like tucidinostat may also result from higher lipophilicity, which can promote off-target or nonspecific effects. Nonetheless, these findings highlight CellNavi’s potential to elucidate the intricate mechanisms of action underlying drug interventions, highlighting an approach to optimize drug efficacy and specificity for targets involving complex downstream signaling pathways.

### CellNavi generalizes to novel cell types

Lastly, we evaluated the generalization capability of CellNavi. We focused on a CRISPR interference screen across HEK293FT and K562 cell lines^67^. The cell types are markedly different in origin and characteristics—HEK293FT cells are derived from human embryonic kidney cells, while K562 cells are derived from human chronic myelogenous leukemia **(Fig. 6a)**. In this experiment, CellNavi was trained on HEK293FT cells, with all K562 cells held out as the test set (**Methods**).

**Figure 6.**
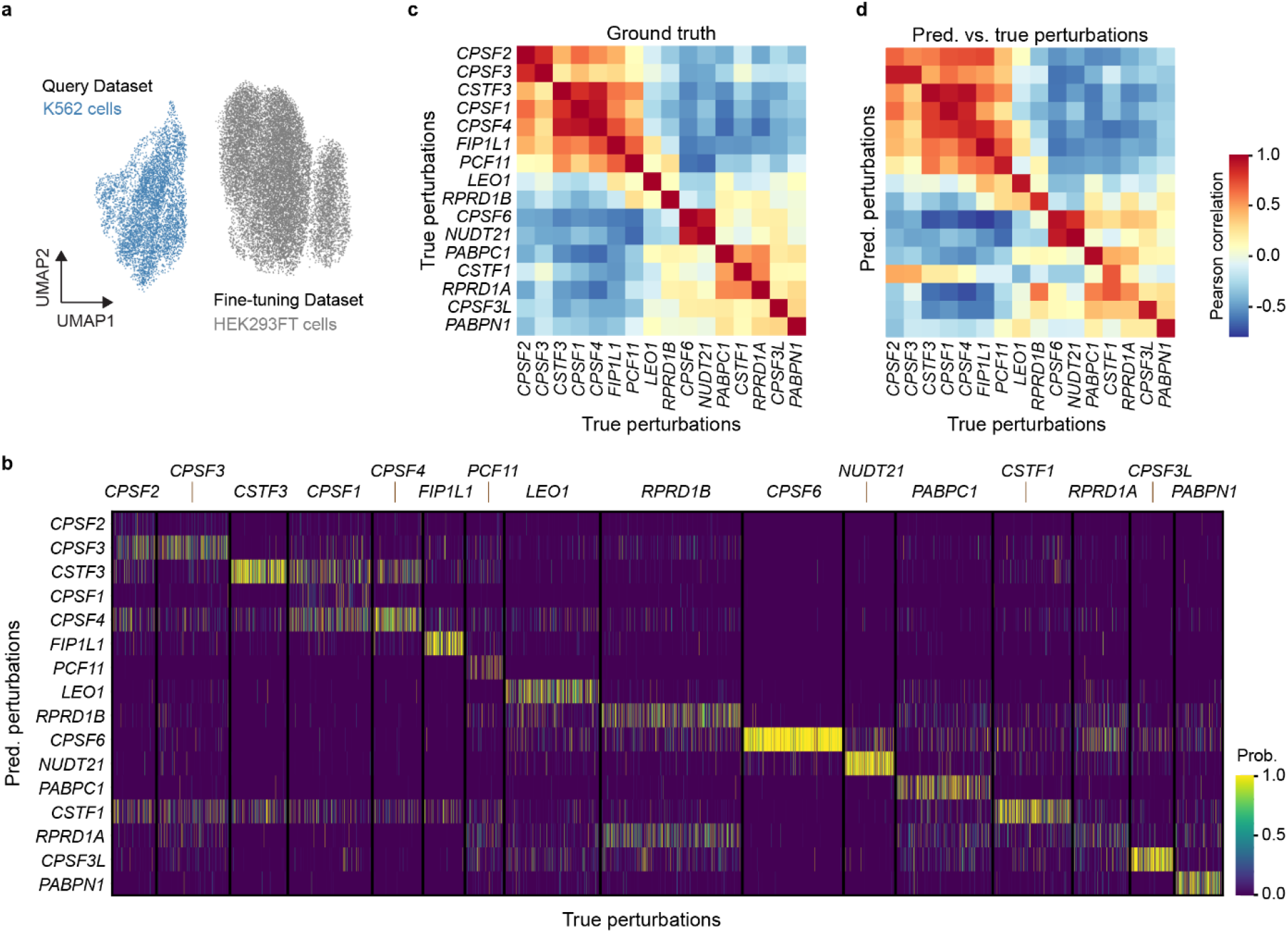
CellNavi predicts driver genes in novel cell types. **a)** UMAP visualization of transcriptomic profiles from Kowalski et al. Gray: HEK293FT cells used for model fine-tuning. Blue: K562 cells for model test. **b)** Predicted perturbations vs. True perturbations in K562 cells. Each row represents a predicted perturbation, and each column represents a cell, whose true perturbation is labeled on top. **c)** Heatmap showing average Pearson correlations over transcriptomic profiles between each pair of perturbations in K562 cells. **d)** Heatmap showing average Pearson correlations over transcriptomic profiles between predicted perturbations and true perturbations in K562 cells. Row: predicted perturbations. Column: true perturbations.

For the 16 perturbations targeting the cleavage and polyadenylation (CPA) regulatory machinery (**Fig. 6a**), CellNavi achieved a macro F1 score of 0.432 on top-1 predictions (**Fig. 6b**). The model misclassified some genes encoding components of the CPSF and CSTF complexes, likely due to their similar post-perturbation transcriptomic profiles (**Fig. 6c**). However, the model performed well in predicting *CPSF6* and *NUDT21*, which exhibit highly similar transcriptomic profiles post-perturbation. Interestingly, despite distinct post-perturbation transcriptomic profiles for *RPRD1A* and *RPRD1B* perturbations, the model confused these genes in many cases. As the protein products of these genes form heterodimers to dephosphorylate the RNA polymerase II C-terminal domain^68^, the model may be prioritizing functional interactions and shared pathways over expression differences, leading to the misinterpretation of these genes.

By comparing the similarities between cell groups stratified by true versus predicted perturbations, we found that both intra- and inter-perturbation correlations for predicted labels closely mirrored those of the true labels **(Figs. 6c–d and Supplementary Fig. 10)**. This suggests that cells grouped by predicted perturbations exhibit gene expression signatures highly similar to those grouped by true perturbations. While prediction accuracy may partly benefit from conserved perturbation effects across cell types, CellNavi remains effective even when applied to cell types markedly different from those used in training, demonstrating robust generalization across diverse cellular contexts.

## Discussion

Understanding the regulatory mechanisms that govern cell identity and transitions stand a central challenge in cell biologyL6,69–72. In this study, we introduce CellNavi, a deep learning framework designed to identify driver genes—key factors that orchestrate complex cellular transitions—across diverse biological contexts. By modeling cell states on a biologically informed manifold constructed from large-scale single-cell transcriptomic data and gene graph priors, CellNavi achieves accurate and generalizable predictions across multiple tasks and datasets.

Describing cell states on a manifold that captures their biological dimensions has been a long-lasting endeavor^32,73–76^. Here, we utilized a structured gene graph derived from NicheNet to facilitate cell state manifold learning via deep neural networks. NicheNet is a comprehensive gene-gene graph integrating both gene regulatory networks and intercellular signaling pathways. This prior improved the accuracy for driver gene prediction compared to alternative or randomized graphs **(Extended Data Table 1)**. Also, integrating prior gene graphs allowed CellNavi to place greater emphasis on transcription factors (TFs), which are crucial for defining cell states and orchestrating transitions^3,10,69^ (**Supplementary Figs. 11-12**). This explicit focus on regulatory elements provides CellNavi with a distinct advantage to model complex biological processes and highlights the value of graph-based learning in improving model interpretability and biological relevance. However, we caution that attention mechanisms do not equate to mechanistic interpretability. The explainability remains a critical challenge for deep learning models, including CellNavi. Future work should develop tools to visualize and interpret how graph structures and attention dynamics shape predictions of driver genes.

Our construction of cell-type-specific graphs involves removing edges for genes with zero expression, based on a simplified assumption that such genes are unlikely to participate in active regulation. Consistent with the previous practices in single-cell foundation models^12,13,17^ and cell-type-specific protein representation^77^ learning, we expect this filtering to help reduce noise and highlight biologically relevant interactions. Yet, we recognize that zero expression values may also stem from technical artifacts such as dropout or low sequencing depth, rather than true biological absence. Future studies should assess alternative strategies, such as imputation or single-cell-level network construction^78^, to balance denoising and information retention.

Inherent noise in biological data presents a significant challenge for modeling. To mitigate technical variability, such as dropout events and differences in sequencing depth, we employed a down-sampling recovery pretraining strategy with a mixed down-sampling rate. This strategy aligns input data of varying depths and improves robustness in handling real-world datasets. Additional noise arises from variability in CRISPR perturbation efficiency, including fluctuating perturbation success rates and off-target effects caused by intrinsic cellular stochasticity. Although CRISPR screens provide a rich and diverse dataset for CellNavi training, this noise may lead to inconsistent labels and biased learning. To mitigate this, future efforts could pool data from multiple batches, sources, and sgRNAs to reduce biases associated with specific experimental conditions. Additionally, integrating orthogonal perturbation data, such as chemical treatments, could complement CRISPR-based data and further enhance model robustness.

CellNavi represents a pioneering effort to benchmark the performance and generalization capacity of deep learning methods on driver gene identification task. While the results are promising, several limitations remain. First, the current pipeline requires fine-tuning on single-cell CRISPR screen data relevant to the system of interest. While our proof-of-concept test involving HEK293FT and K562 cells demonstrated promising results **(Figure 6)**, the extent to which CellNavi can generalize to entirely new cell types or experimental systems remains unclear. Addressing this will require testing across more diverse contexts and quantifying the “distance” between systems to determine when fine-tuning is necessary. A long-term goal is to reduce dependence on such datasets by developing models that generalize with minimal experimental effort.

Second, CellNavi cannot yet generalize to novel genes, which limits its broader applicability. Expanding this capacity would require capturing gene networks and representations that enable extrapolation beyond the training dataset. While single-cell CRISPR experiments encompassing a broader range of target genes and cell types are desirable, integrating generative models to infer missing relationships could further improve the model’s capacity to handle novel genes.

Third, CellNavi lacks the ability to accurately model long-range transitions due to its reliance on CRISPR perturbations and static snapshots of transcriptomic data. Many biological processes, such as differentiation and disease progression, unfold gradually through transient states not captured in steady-state data. Incorporating time-resolved single-cell data measurements could help construct dynamic manifolds that better reflect these processes.

Despite these challenges, CellNavi marks a significant advance in modeling cell state transitions and identifying their genetic drivers. By combining biologically informed priors with advanced deep learning techniques, CellNavi achieves high accuracy and generalizability in diverse biological contexts. As we continue to refine and expand models like CellNavi, we are paving the way for novel treatments targeting the root causes of diseases with unprecedented specificity.

## Acknowledgements

We thank Dr. Zeyu Chen, Dr. Sida Shao, Dr. Chuan Cao and Zeyu Tang for constructive suggestions and feedback on the manuscript and Dr. Tie-Yan Liu for the supervision. We thank Jingyun Bai for graphic designs and illustrations.

## Author Contributions

Conceptualization, P.D., T.W., S.Z., and C.L.; Data curation, T.W. and P.D.; Methodology, Y.P., F.J., T.W., S.Z., C.L., and P.D.; Model implementation – pre-train, F.J., Y.P., G.L., and H.X.; Model implementation – fine-tuning and inference, Y.P., F.J., T.W., C.L., and Q.J.; result analysis, T.W., Y.P., F.J., P.D., Y.M., and X.L., result interpretation, T.W., P.D., Y.P., F.J., Y.M.; Writing-original draft, P.D., T.W., C.L., S.Z., F.J., Y.M., and Y.P.; Writing-revision, P.D., C.L., S.Z., H.L., Y.P., H.X.; Supervision, H.L.. All authors have read and approved the manuscript.

## Methods

### Input embeddings

In CellNavi, we use single-cell raw count matrices as the only input. Specifically, single-cell sequencing data is processed into a cell-gene count matrix,**X** ∈ ℝ^*N*×*G*^, where the each element **X**_*n,g*_ represents the expression of the *n*-th cell and the *g*-th gene (or read count of the *g*-th RNA).

To better describe a gene’s state in a cell, we involve both gene name and gene expression information in its input embeddings. Formally, the input embedding of a token is the concatenation of gene name embedding and gene expression embedding.

#### Gene name embedding

We use a learnable gene name embedding in CellNavi. The vocabulary of genes is obtained by taking the union set of gene names among all datasets. Then the integer identifier of each gene in the vocabulary is fed into an embedding layer to obtain its gene name embedding. Additionally, we incorporate a special token CLS in the vocabulary for aggregating all genes into a cell representation. The gene name embedding of cell *n* can be represented as 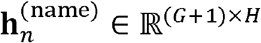:

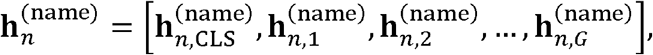

Where *H* is the dimension of embeddings, which is set to 256.

#### Gene expression embedding

One major challenge in modeling gene expression is the variability in absolute magnitudes across different sequencing protocols^13^. We tackled this challenge by normalizing the raw count expression for each cell using the shifted logarithm, which is defined as,

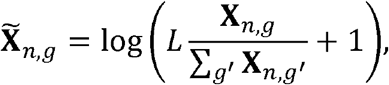

where **X**_*n,g*_ is the raw count of gene *g* in cell *n, L* is a scaling factor and we used a fixed value *L* = 1 × 10^4^ in this study, and 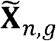 denotes the normalized count. Finally, a linear layer was applied on the normalized expression 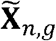 to obtain the gene expression embedding. For the CLS token, we set it as a unique value for gene expression embedding. The gene expression embedding of cell *n* can be represented as 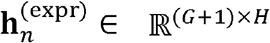:

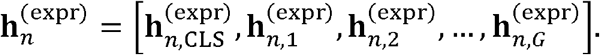

The final embedding of cell *n* is defined as the concatenation of 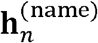 and 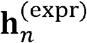:

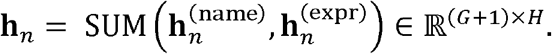

### Cell Manifold Model

#### Model architecture

The Cell Manifold Model, is composed of 6 layers of a Transformer variant that is designed specifically for processing graph-structured data (GeneGraph Attention Layers)^19^. The encoder takes the input embeddings to generate cell representations and uses only genes with non-zero expressions. To further speed up training, also as an approach of data augmentation, we performed a gene sampling strategy by randomly selecting at most 2,048 genes as input. It should be noted that the strategy is only applied during training, all non-zero genes are included at inference stage to avoid information loss. We use 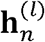 to represent the embedding of cell *n* at the *l*-th layer, where 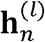 is defined as:

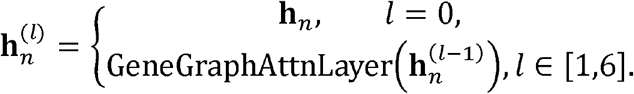

The multi-head attention module in each GeneGraph Attention Layer consists of 3 components. In addition to a self-attention module, a centrality encoding module and a spatial encoding module are also incorporated to modify the standard self-attention module for graph data integration.

We start by introducing the standard self-attention module. Let *N*_head_ be the number of heads in the self-attention module. In the *l*-th layer, *i*-th head, self-attention is calculated as:

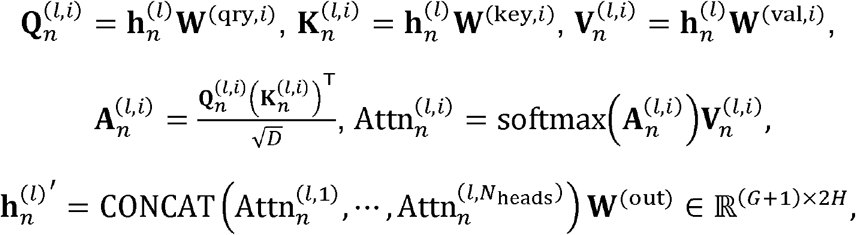

where **W**^(qry,*i*)^, **W**^(key,*i*)^, **W**^(val,*i*)^ ∈ ℝ^2*H*×*D*^ are learnable matrices that project input embedding 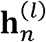 of cell *n* in to 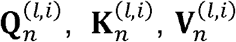, the symbol 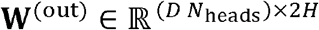 is a learnable linear projection that refines the output of multi-head attention, and *D* is the feature dimension for each attention head which satisfies *D N*_heads_ = 2*H*. The output of multi-head attention 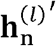 is then passed through a layer normalization layer and a multi-layer perceptron model (MLP), producing the final output 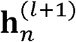 as the input to the next layer.

The standard attention mechanism only processes features on each single gene, while the gene graph provides the relational information among the genes. To incorporate the gene graph information into the model, the centrality encoding module projects the relational information into the regulatory activity feature of each single gene, and the spatial encoding module directly incorporates the relational information with the attention mechanism. To be more specific, we define 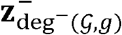 and 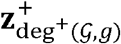, learnable embeddings describing in-degree deg^−^ and out-degree deg^(^ of gene *g* on the gene graph 𝒢. We add these embeddings to the gene embeddings to update cell encoding:

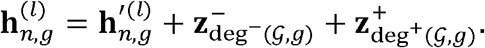

This cell encoding update by the centrality encoding module is applied before the self-attention module.

The spatial encoding module aims to capture regulation relations between genes from the gene graph. For this purpose, we generate the distance matrix **S** ∈ ℕ^*G*×*G*^ containing shortest distances between gene pairs on the gene graph 𝒢. We assign each element in **S** as a learnable bias added to attention weights:

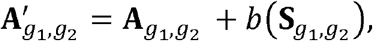

Where *b* is a learnable scalar-valued function of the distance 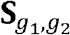. It assigns a special value to genes that are not connected to the graph. We use **A**′, in place of the original attention weights **A** in the standard self-attention module when computing self-attention in our model. In our implementation, we apply layer normalization and an MLP before computing multi-head self-attention. The cell representation output from the CMM, 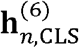, is subsequently passed through a fully connected layer, where the dimensionality is increased from 256 to 2,048. This resulting value serves as the cell coordinate for cell *n*, denoted as **CDR**_*n*_.

#### CMM Pretraining Task

The CMM is expected to generate cell coordinates that parameterize the intrinsic features/variables (that are much less than the dimensions in the raw gene expression profile representation) of a cell state and maintain cell similarity in the vector space, so as to provide a concise and biologically relevant representation for the DGP to consume. To achieve this, we design a down-sampling reconstruction pretraining task, which asks the CMM to produce a cell coordinates of a down-sampled gene expression 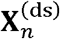 of a cell *n*, that allows a separate decoder model to reconstruct the original gene expression **X**_*n*_ of that cell as accurate as possible. In order to achieve this, the CMM is enforced to capture the co-varying patterns among the raw gene expression dimensions, hence helping the CMM to extract the underlying intrinsic variables.

Specifically, for the down-sampling process, we down-sample the raw count expression of each gene via a binomial distribution. The down-sampled expression 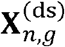 of the *n*-th cell and the *g*-th gene is produced by:

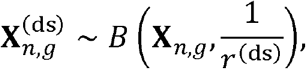

Where **X**_*n,g*_ is the raw count of gene *g* in cell *n, r*(ds) is the down-sample rate which is uniformly sampled from [1,20), and *B* denotes the binomial distribution. The decoder is an MLP consisting of two linear layers. For each down-sampled gene expression, the decoder concatenates the cell coordinates **CRD**_*n*_ of 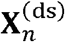 produced by the CMM and the embedding of that gene as the direct input to the MLP. The MLP output comes in the same shape as **X**_*n*_.

The learning objective for reconstructing the original gene expression profile **X**_*n*_ from the down-sampled version 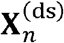 is:

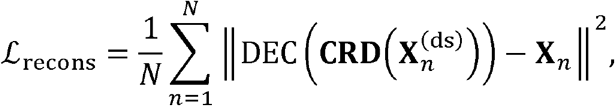

where ‖ · ‖^2^ represents the squared 2-norm of a vector. Both the CMM and the decoder are optimized. After pretraining, the CMM is to be used for driver gene prediction, while the decoder is discarded.

### Driver Gene Predictor

The driver gene classifier is an MLP consisting of two linear layers. It is optimized to predict the perturbed genes from a pair of cell coordinates output by the Cell Manifold Model. To be more specific, transcriptomes of source cell x_src_ and target cell x_tgt_ are mapped to cell coordinates **CRD**_src_ and **CRD**_tgt_ with the Cell Manifold Model. For the direct input features, the DGP concatenates the two cell coordinates and then proceeds with an MLP, which outputs the logits of genes. We use the cross-entropy loss for training the DGP:

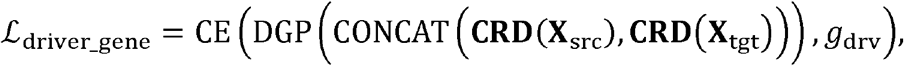

where 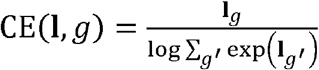 is the cross-entropy loss, and *g*_drv_ denotes the driver gene corresponding to **x**_src_ and **x**_tgt_. The loss is finally averaged over all (**x**_src_, **x**_tgt_, *g*_crv_) tuples in the dataset. The pretrained CMM used to produce **CRD**_src_ and **CRD**_tgt_ is also finetuned together with the DGP by this loss.

Additional training details for CellNavi are available in **Supplementary Note 4**.

### Baselines

#### SCENIC / SCENIC

For each test dataset, SCENIC+ inferred a gene regulatory network (GRN), identified regulons 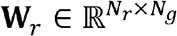, and computed regulon activity 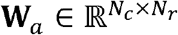 in regulatory network the cells, where N_*r*_,*N*_*g*_,*N*_*c*_ represent the number of identified regulons, genes, and cells in the test dataset, respectively. **w**_*r*_ is a learnt matrix containing the weights of genes for different regulons, and **w**_*a*_ indicates the regulon activities for each cell. Then we used **w**_*g*_ = **w**_*a*_ **w**_*r*_ to represent the regulatory importance of each gene in cells. Then, genes in each cell were ranked based on the value of their regulatory importance (elements in **w**_*r*_), as high value indicates a possible role in controlling cellular identity. We applied SCENIC+ to Norman et al. and Schmidt et al. datasets. Only genes appeared in perturbation pools in these datasets participated in the ranking based on w_*g*_. Hyperparameters of GRN inference, regulon identification, and regulon activation were set to default. Cells with no regulon activated were removed from our analysis. SCENIC+ analysis was realized by `pyscenic 0.12.1`.

#### Other GRNs

We constructed gene regulatory networks (GRNs) using three alternative methods: GRNBoost2, GENIE3, and RENGE, following default parameters from prior studies where applicable. Due to computational memory constraints, we limited the analysis for GENIE3 and RENGE to the top 5,000 highly variable genes. For GENIE3 and GRNBoost2, we utilized the SCENIC implementation to infer GRNs. For RENGE, which is designed to infer GRNs using time-series scRNA-seq data, we adapted the method to work with static single-cell RNA-seq data. After constructing GRNs with these methods, we applied the same downstream analysis protocol as described for the SCENIC pipeline.

#### In silico perturbation

In silico perturbation (ISP) methods, such as GEARS, are capable of predicting transcriptomic outcomes of genetic perturbations. We trained GEARS model on the datasets mentioned in the corresponding tasks. For evaluation, we computed the cosine similarity between predicted transcriptomic profiles upon different perturbations, and that of cells from test datasets. Driver genes were predicted based on the similarities, and high values in similarity indicate the potential to be driver genes. GEARS analysis was realized by cell-gears 0.1.1. Data processing and training followed the data processing tutorial (https://github.com/snap-stanford/GEARS/blob/master/demo/data_tutorial.ipynb) and training tutorials (https://github.com/snap-stanford/GEARS/blob/master/demo/model_tutorial.ipynb).

#### DGE analysis

Differential gene expression (DGE) analysis is the most frequently used method to reveal cell-type specific transcriptomic signature. Initially, cells from the test datasets were normalized and subjected to logarithmic transformation. Subsequently, we applied the Leiden algorithm, an unsupervised clustering method, to categorize the target cells into distinct groups. The number of clusters for each test dataset was set to range from 20 to 40, ensuring that cellular heterogeneity was maintained while providing a sufficient number of cells in each group for robust statistical analysis. We selected source cells to serve as a reference for comparison and performed DGE analysis on each target cell group against this reference. The Wilcoxon signed-rank test was employed to determine statistical significance. Then, significant genes were ranked according to their log-fold changes in expression as potential driver genes. Both unsupervised clustering and DGE analysis were conducted using the package `scanpy 1.9.6`.

### The prior gene graph

The prior gene graph was constructed from NicheNet, where gene regulatory network and cellular signaling network were integrated. The gene graph is a directional graph. To be more specific, for each gene node on the graph, the number of incoming edges corresponds to the genes that regulate it, while the number of outgoing edges represents the genes it regulates. In our approach, a connection was established between two genes if they were linked in either of the individual networks. The resulting integrated graph features 33,354 genes, each represented by a unique human gene symbol, and includes 8,452,360 edges that signify the potential interactions. The unweighted versions of NicheNet networks were used in our approach. For each cell, we remove the gene nodes with values of 0 in the raw count matrix of the single-cell transcriptomic profile, to construct the cell type-specific gene graph. During pre-training, when down-sampling is performed on single-cell transcriptomes, only the non-zero genes included as model input are retained to generate sample-specific graphs that guide the model’s task.

To evaluate the impact of graph connectivity and structure, we generated alternative graph configurations as follows:

1. **Fully Connected Graph**: A maximally connected graph where every pair of genes is connected by an edge of equal weight.
2. **Sparsified Graphs**: Graphs were created by down-sampling the total number of edges from the original graph to 1/10 and 1/20 of its total edges, enabling an evaluation of how reduced connectivity affects performance.
3. **Random Graphs**: Randomized graphs were generated while preserving the number of nodes and certain structural properties of the original graph, such as self-loops. Edges were introduced probabilistically to maintain overall consistency with the original graph’s sparsity and connectivity.

### Datasets

#### Human cell atlas (HCA)

We downloaded all single-cell and single-nucleus datasets sourced from contributors or DCP/2 analysis in *Homo sapiens* up to March 2023, accumulating approximately 1.5 TB of raw data. We retained all experiments that included raw count matrices and standardized the variables to gene names using a mapping list obtained from Ensembl (https://www.ensembl.org/biomart/martview/574df5074dc07f2ee092b52c276ca4fc).

#### Norman et al

This dataset (GEO: GSE133344) measures transcriptomic consequences of CRISPR-mediated gene activation perturbations in K562 cell line. We filtered this dataset, removing cells whose counts number is lower than 3500. After filtering, this dataset contains 105 perturbations targeting different genes, and 131 double perturbations targeting two genes simultaneously. We used unperturbed cells (with non-targeting guide RNA) as source cells and perturbed cells as target cells.

#### Schmidt et al

This dataset (GEO: GSE190604) measures the effects of CRISPR-mediated activation perturbations in human primary T cells under both stimulated and resting conditions. For our analysis, we excluded cells not mentioned in metadata and removed genes appeared in less than 50 cells. Gene expression levels of single-guide RNA were deleted in order to avoid data leakage. We used unperturbed cells as source cells and perturbed cells as target cells. We also excluded cells without significant changes after perturbation following the procedure proposed by Mixscape tutorial via package pertpy. Default parameters were used for Mixscape analysis.

#### Cano-Gamez et al

This dataset (EGA: EGAS00001003215) comprises naïve and memory T cells induced by several sets of cytokines. With cytokine stimulation, T cells are expected to differentiate into different subtypes. We took cells not treated by cytokines as source cells and cytokine-stimulated cells as target cells. This experimental set reflects the differential process of human T cells. For our analysis, clusters 14-17 were excluded as their source cell could hardly be decided.

#### Fernandes et al

This dataset comprises heterogeneous dopamine neurons derived from human iPSC. These neurons were exposed to oxidative stress and ER stress, representing PD-like phenotypes. We followed preprocessing procedures as mentioned in the original GitHub repo (https://github.com/metzakopian-lab/DNscRNAseq/blob/master/preprocessing.ipynb).

#### Tian et al

This dataset comprises iPSC-derived neurons perturbed by more than 180 genes related to neurodegenerative diseases. CRISPR interference experiments with single-cell transcriptomic readouts were conducted by CROP-seq. For our analysis, we removed genes appeared in less than 50 cells. We used unperturbed cells as source cells and perturbed cells as target cells.

#### Srivatsan et al

This dataset (GEO: GSE139944) contains transcriptomic profiles of human cell lines perturbed by compounds. For our study, we utilized K562 cell line cells perturbed by HDAC inhibitors. We used unperturbed cells as source cells, and chemically perturbed cells as target cells. This set represents the process of cellular transition caused by drugs.

#### Kowalski et al

This dataset (GEO: GSE269600) measures the transcriptional consequences of CRISPR-mediated perturbations in HEK293FT and K562 cells. For our analysis, we excluded perturbations which consist of less than 200 cells. Cells with minimal perturbation effects were removed from downstream analysis. We used cells from control groups as source cells, and perturbed cells as target cells.

### T cell differentiation analysis

#### Identification of cellular phenotypic shift

We computed ‘transition score’ to identify cellular phenotypic shift on the transcriptomic level. We selected canonical marker genes associated with IFNG and IL2 secretion, and Th2 differentiation. Then, we computed transition score based on the mean expression level of these marker genes. That is: 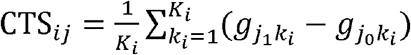, where CTS_ij_ is the transition score of phenotypic shift type _*i*_ in source-target cell pair 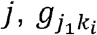 and 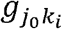 are normalized gene expression levels of marker gene *k* for cell-state transition type *i* in target cell *j*_1_ and its source cell *j*_0_. The total number of maker genes for phenotypic shift type *i* is represented with *K*_*i*_. Then, classes of phenotypic changes were annotated based on transition score. Transition scores are calculated via function `tl.score_genes` from `scanpy` package with default parameters.

#### Cell type classification with predicted driver genes

We selected a series of genes related to transition mentioned above from previous studies (**Supplementary Table 4**). We used the term ‘likelihood scores’ to describe the probability of a gene to be a driver factor predicted by the model, that is:

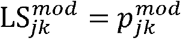

where 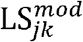 means the likelihood score for gene *k* in source-target cell pair *j* from model *mod*, and 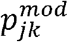 represents the probability predicted by model *mod*. For our analysis, the *mod* could be CellNavi, or baseline models.

Then, we aggregate likelihood scores into ‘prediction scores’ to evaluate the performance of different models:

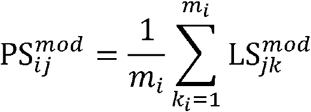

where 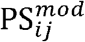 is the prediction score of cell-state transition type *i* in source-target cell pair *j* predicted by model *mod*. The number of candidate driver genes for each phenotypic changing type *i* is *m*_*i*_. Ideal prediction should reflect similar patterns as shown by cellular transition score mentioned above. To evaluate it quantitatively, we trained decision tree classifiers with prediction scores as input to test whether predictions scores would faithfully demonstrate cell-state transition types. Classifiers were trained for each method independently, and 10-fold cross-validation was conducted. Classifiers were implemented via shallow decision trees using the `sklearn` package.

#### Gene Ontology enrichment analysis

We used gene ontology (GO) enrichment analysis to explore drugs’ mechanism of actions. For each drug compound, the top 50 genes with highest scores predicted by CellNavi were used for GO enrichment analysis. The significant level was chosen to be 0.05, and the Benjamini-Hochberg procedure was used to control the false discovery rate. For implementation, we used package `goatools` for GO enrichment analysis.

#### Molecular docking

We performed molecular docking for panobinostat and tucidinostat, with a reference protein structure obtained from the PDB entry 3MAX. The ligand structures from PDB entries 3MAX and 5G3W were used to guide the initial placement of panobinostat and tucidinostat, ensuring the pose correctness of the warheads and major scaffolds. Based on such initial poses, local optimizations were performed with AutoDock Vina. PyMol was used for structure visualization.

## Data Availability

HCA data for Cell Manifold Model training is downloaded from the HCA data explorer (https://explore.data.humancellatlas.org/projects). The Norman et al.^32^, Tian et al.^52^, and Srivatsan et al.^58^ datasets were downloaded from the scPerturb project ^81^ on Zenodo (https://zenodo.org/records/10044268). The raw count data of Schmidt et al.^30^ dataset was downloaded from the National Institutes of Health GEO with accession number GSE190604, and its metadata was downloaded from Zenodo (https://zenodo.org/records/5784651). The Cano-Gamez et al.^38^ dataset was downloaded from the Open Target Platform of this project (https://www.opentargets.org/projects/effectorness). The Fernandes et al.^50^ dataset was downloaded from ArrayExpress with accession number E-MTAB-9154. The preprocessed Kowalski et al^67^ dataset was downloaded was downloaded from Zenodo

(https://zenodo.org/records/7619593#.Y-P7Zi1h2X0). For trajectory reconstruction, we used the dataset from GSE132188. For scRNA-seq alignment across varying sequencing depths, we used data from GSE84133, specifically the Human3 sample. PDB entry: 3MAX, 5G3W.

## Code availability

Custom code developed in this study: https://github.com/DLS5-Omics/CellNavi. Additional software packages for modeling and data analysis: python==3.8.19, torch==2.4.0, pandas=2.2.2, numpy==1.23.5, scikit-learn==1.5.1, scipy==1.14.1, networkx==3.3, scanpy==1.10.3. cell-gears==0.1.2, pyscenic==0.12.1, renge==0.0.3, cell-gears=0.1.1, goatools==1.4.12, pertpy (2024.04).

---

**Extended Data Fig. 1.**
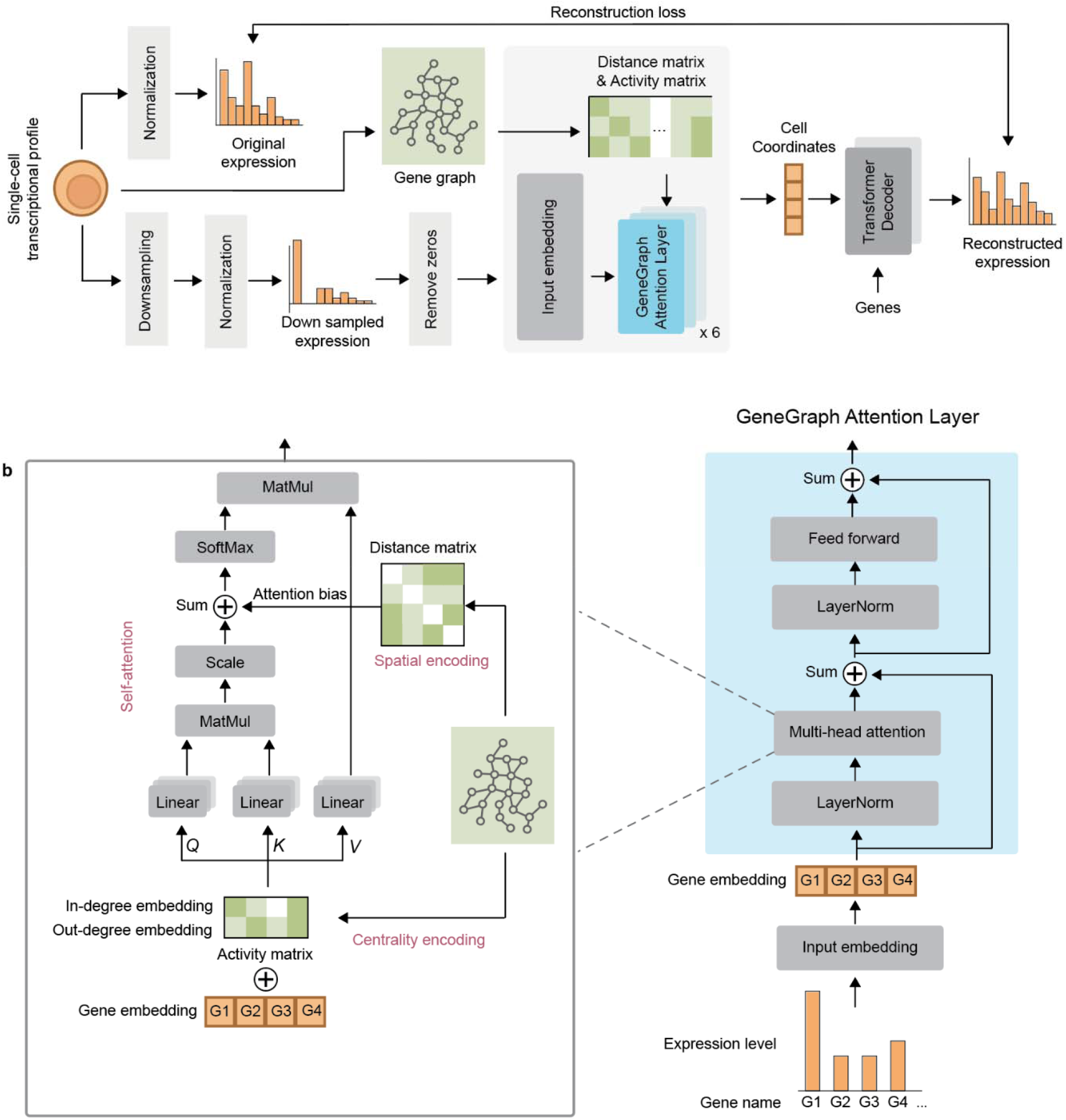
Details of the Cell Manifold Model (CMM) implementation. **a)** The CMM is designed to reconstruct gene expression profiles by leveraging a Transformer variant composed of GeneGraph Attention Layers. **b)** GeneGraph Attention Layer integrates prior gene graph via its multi-head attention layer. To be noted, only sub gene graphs with nodes in the input sample (all non-zero genes) are used for attention calculation.

**Extended Data Fig. 2.**
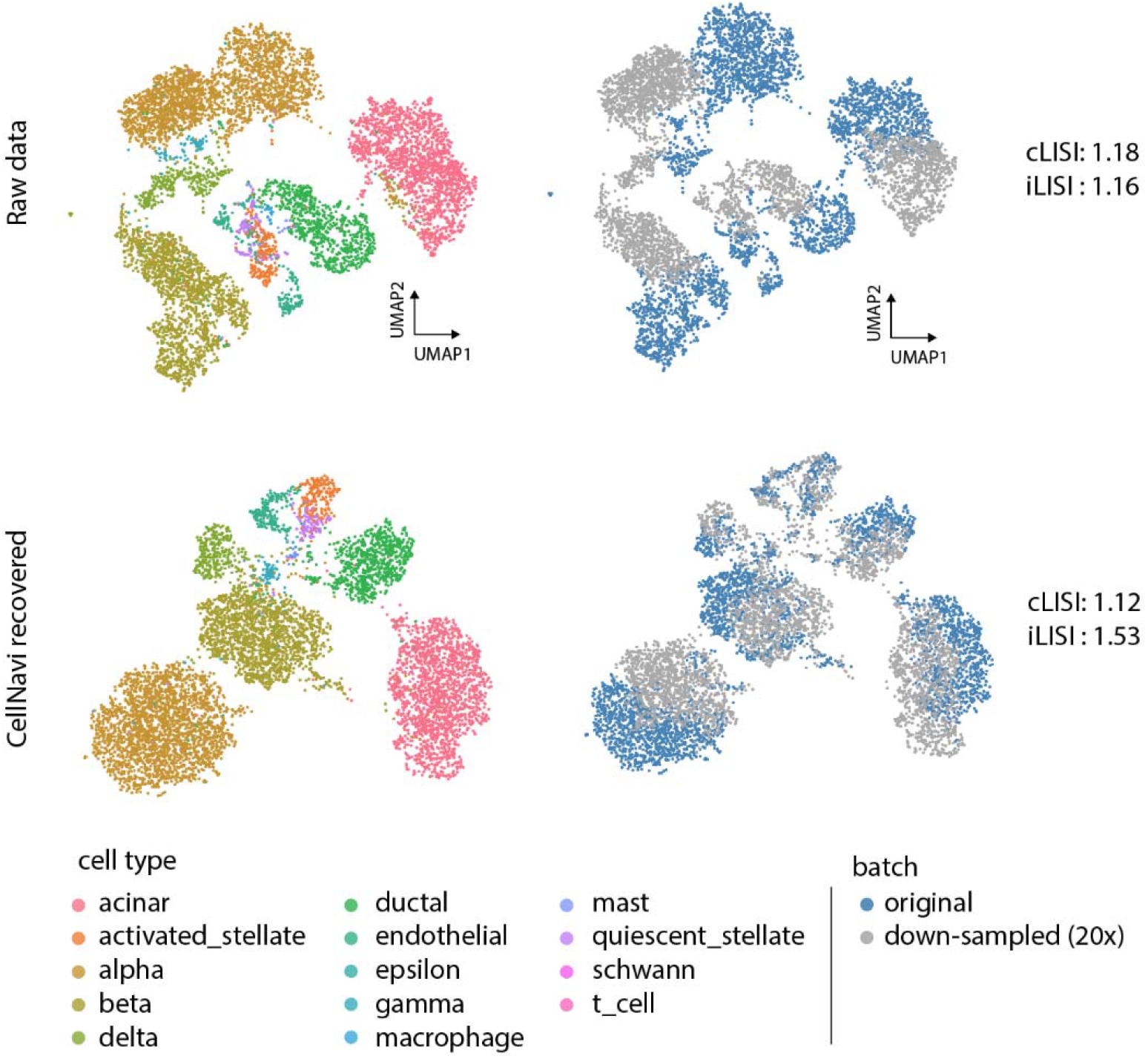
CellNavi can align the same cell types across varying sequencing depths. Upper panel: After 20-fold down-sampling on the expression profile^79^, major cell types remained distinguishable, but a severe batch effect was observed even after using standard integration methods. **Lower panel:** the Cell Manifold Model (CMM) in CellNavi can integrate embeddings from the down-sampled and original profiles, while preserving the biological structure. Integration quality was quantitatively assessed using iLISI (1: worst, 2: best) and cLISI (1: best, 2: worst), which evaluate batch mixing and cell type separation, respectively. See details of experimental rationales in **Supplementary Note 2**.

**Extended Data Fig. 3.**
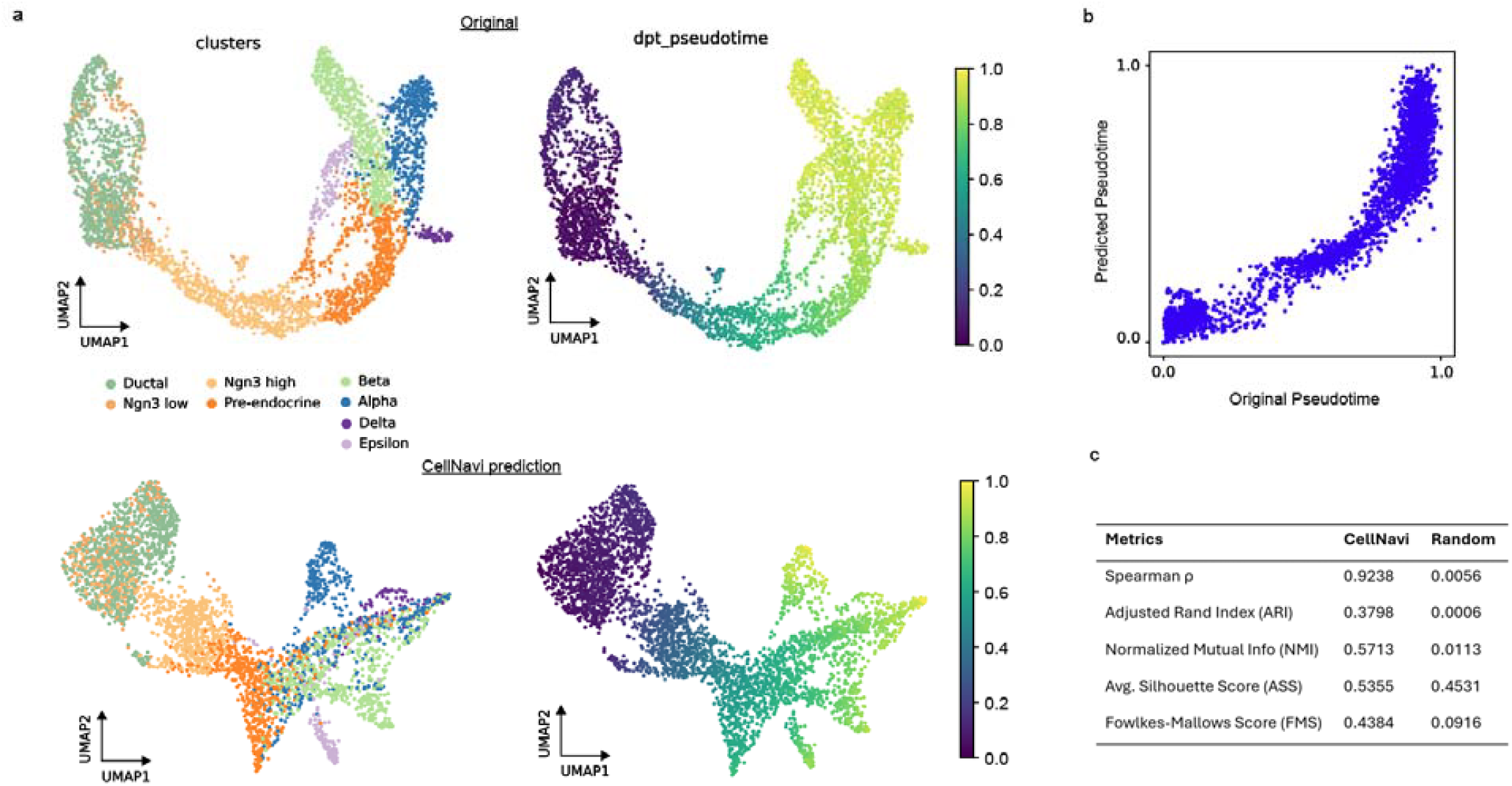
Trajectory reconstruction with the Cell Manifold Model. **(a)** UMAPs colored by cell type (left) and diffusion pseudotime (DPT) (right, indicated by the colorbars), inferred from the original data (upper panel) and CellNavi-predicted data (lower panel). Cell type labels were assigned based on known markers, and pseudotime was computed using Scanpy’s DPT implementation. **(b)** Spearman correlation between pseudotime inferred from the original data (x-axis) and from the CellNavi-predicted data (y-axis) with a quantification in c. **(c)** Summary of quantitative evaluation using Spearman correlation, along with clustering evaluation metrics including Adjusted Rand Index (ARI), Normalized Mutual Information (NMI), Average Silhouette Score (ASS), and Fowlkes-Mallows Score (FMS). Scores computed from random embeddings are included for comparison. We used a mouse pancreatic endocrinogenesis dataset from Bastidas-Ponce et al.^80^. Therefore, we enhanced our model with mouse scRNA-seq for this task.

**Extended Data Fig. 4.**
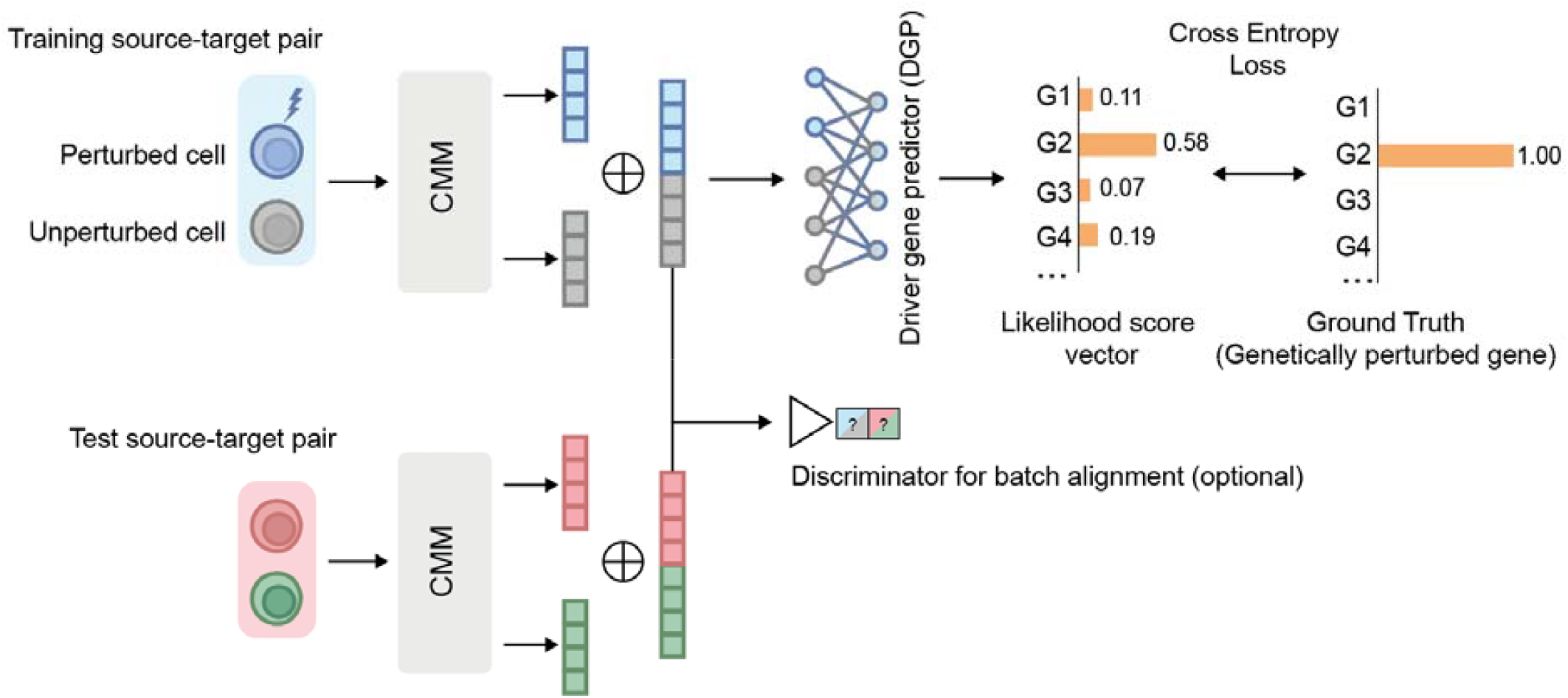
Details of the Driver Gene Predictor (DGP) implementation. The DGP processes concatenated cell coordinates pairs output by the CMM to predict driver genes by generating a likelihood score vector. An optional discriminator is included to align training and test data, ensuring consistency and accuracy in predictions (**Supplementary Note 4**).

**Extended Data Fig. 5.**
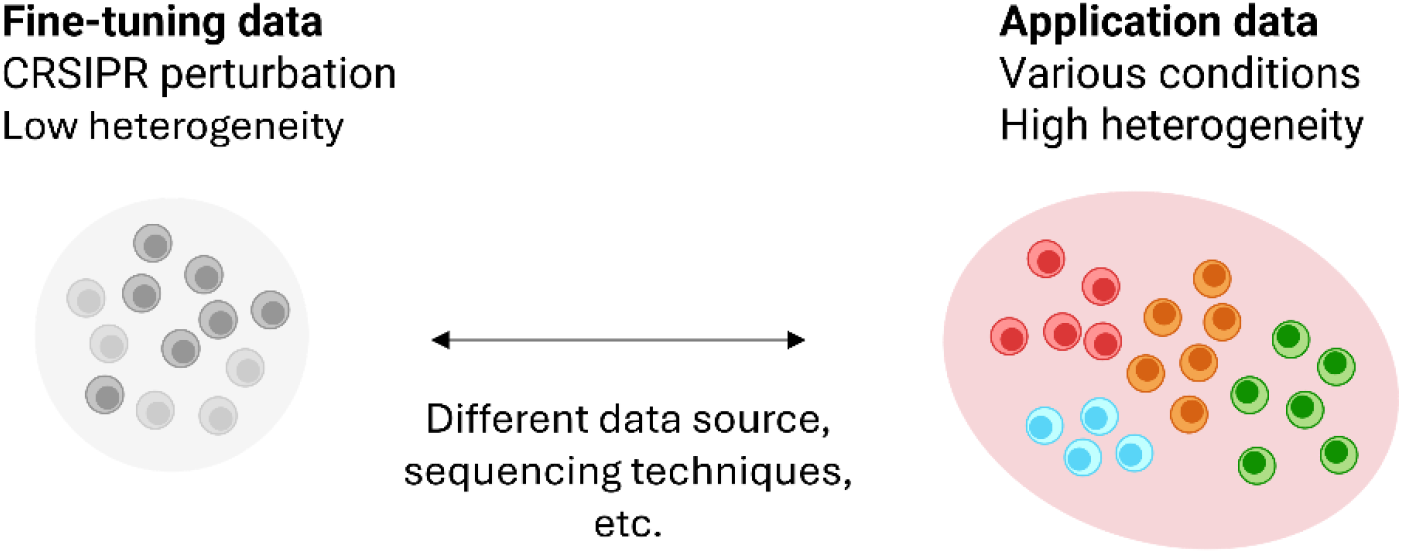
Principle of CellNavi application. The fine-tuning data (left) indicate of CRISPR perturbation datasets with low heterogeneity, typically derived from controlled experiments with limited variability. The application data (right) encompass diverse biological conditions with high heterogeneity, including different cell types, perturbation types, and sequencing platforms. The bidirectional arrow indicates the challenges posed by differences in data sources, sequencing techniques, and biological variability when generalizing the model from fine-tuning to real-world applications. Diagram created with BioRender.com.

**Extended Data Fig. 6.**
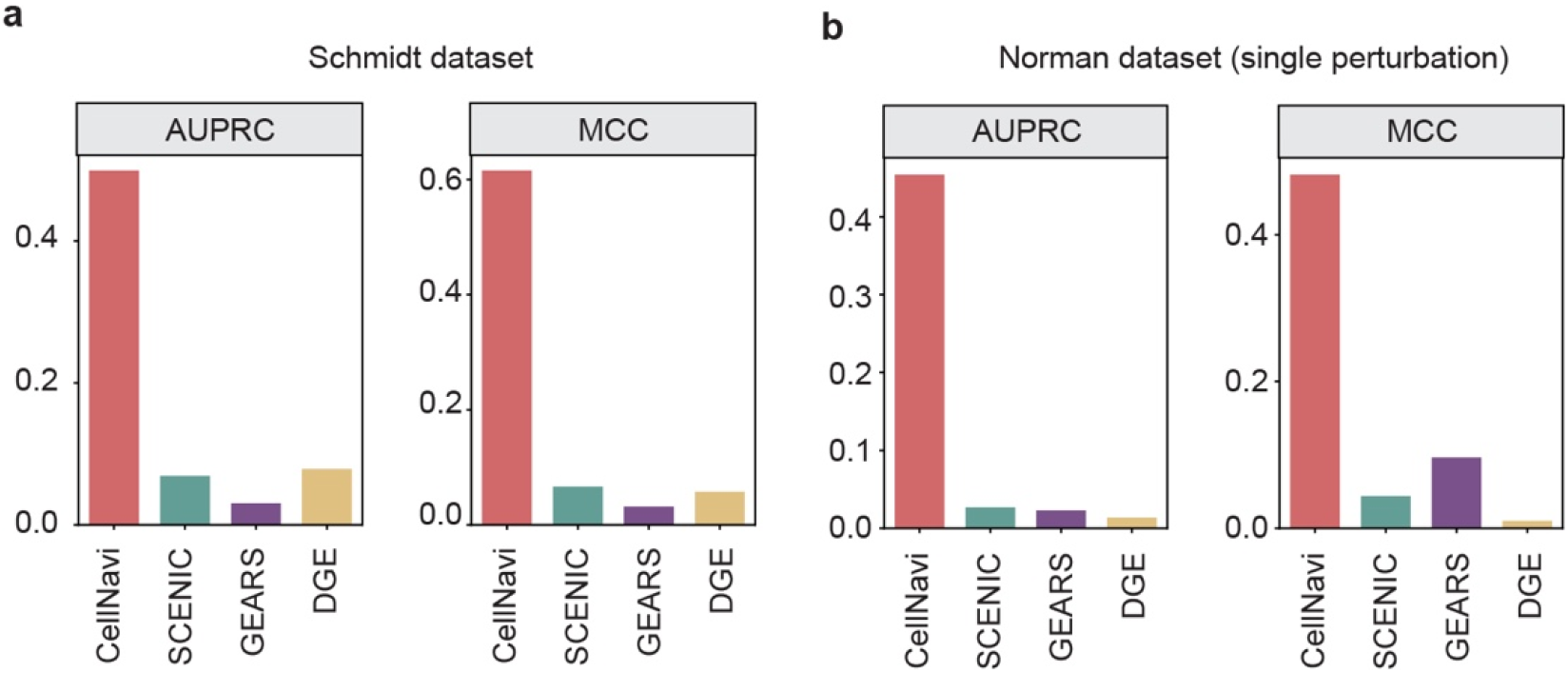
AUPRC (Area Under Precision-Recall Curve) and MCC (Matthews Correlation Coefficient) for driver gene prediction. (a) in the Schmidt dataset and (b) in the Normal dataset (b), comparing CellNavi with alternative methods.

**Extended Data Fig. 7.**
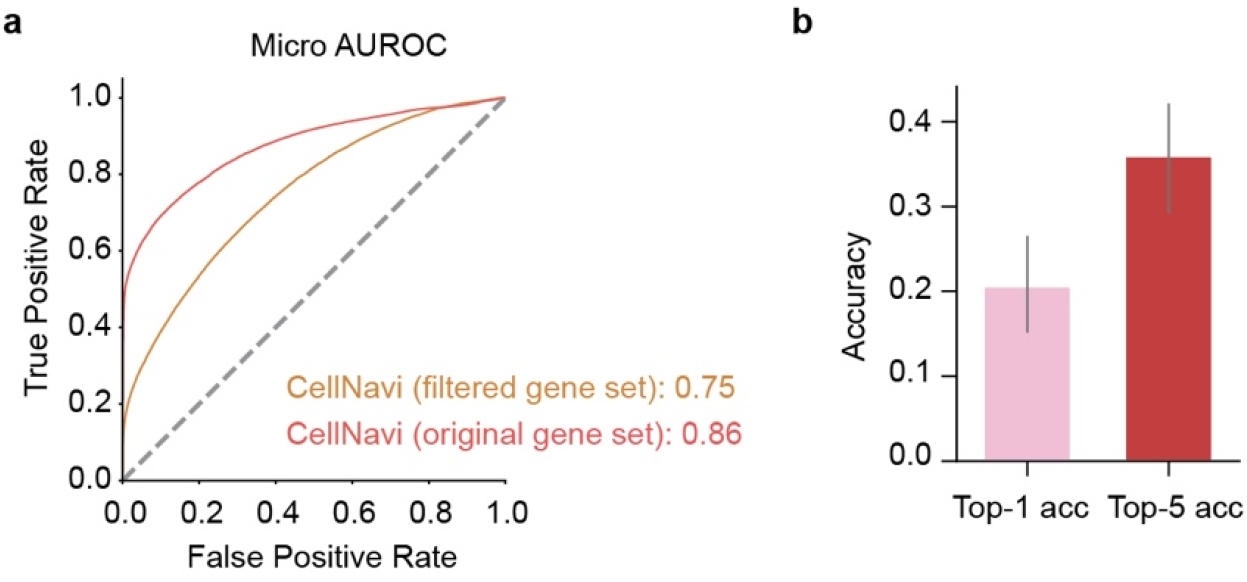
CellNavi identifies driver genes excluded from gene expression profiles. **(a)** Micro-AUROC of CellNavi applied to the original single-cell transcriptomic profile and the filtered transcriptomic profile in which perturbed genes are excluded. **(b)** Top *K* (*K* = 1 or 5) accuracy of CellNavi applied to transcriptomic data with perturbed gene excluded. Error bar, standard error. *n* = 69.

**Extended Data Fig. 8.**
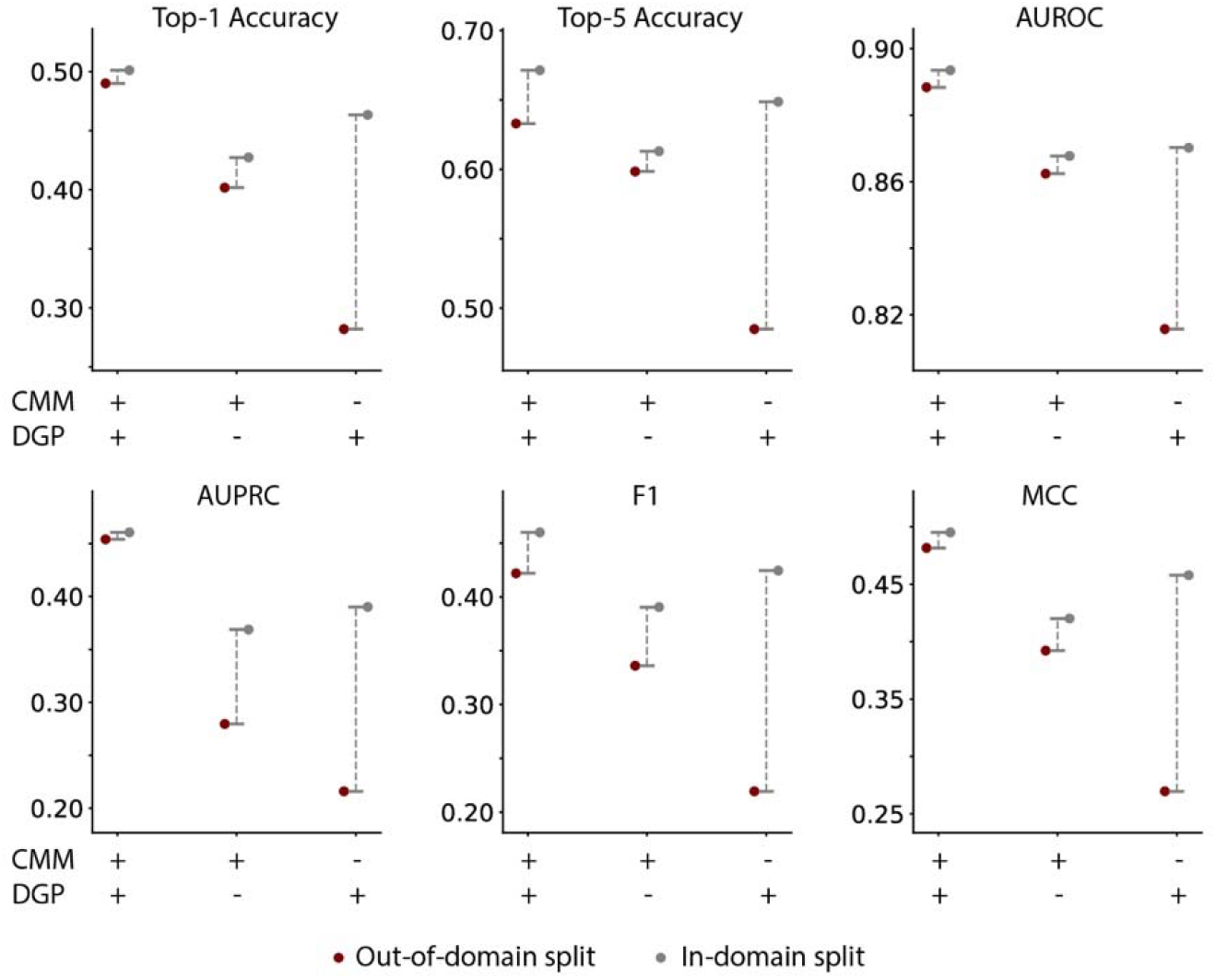
Ablation study on CellNavi components. The figure compares the performance of CellNavi under three configurations: (1) CellNavi with both CMM pretraining and DGP fine-tuning, (2) coupling the DGP with raw gene expression vectors instead of gene embeddings produced by the CMM, and (3) replacing the DGP with a simpler multinomial logistic regression model on top of the CMM. Performance is assessed under two evaluation settings: out-of-domain (the previously used Norman single perturbation split that holds one cluster out from training, red) and in-domain (a random split on the Norman dataset with the same train/test size, gray). Metrics include Top-1 accuracy, Top-5 accuracy, AUROC, AUPRC, F1 score, and MCC (Matthews Correlation Coefficient). Dashed lines indicate performance change between evaluation settings.

**Extended Data Table 1.**
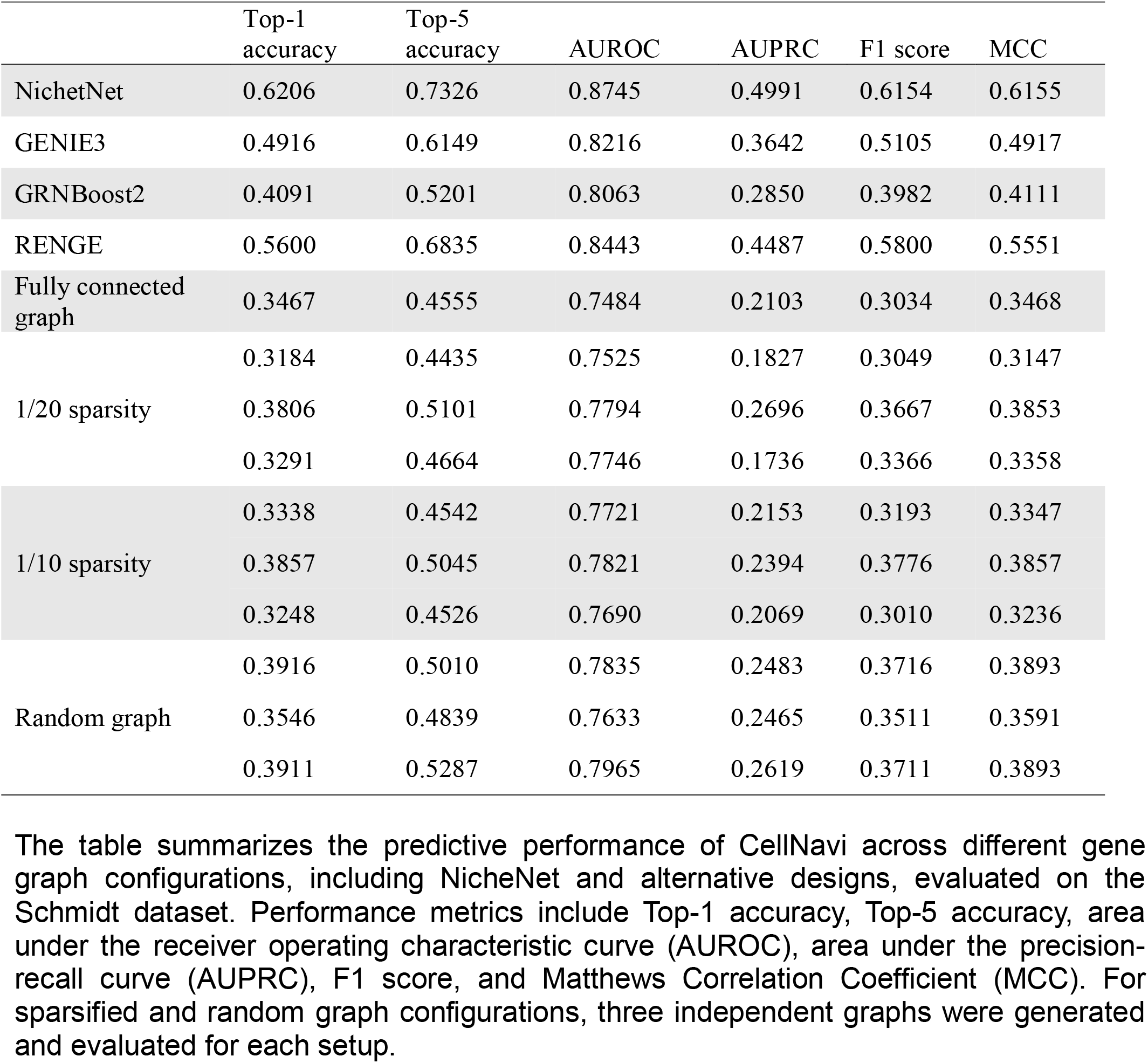
Comparison of the predictive performance of CellNavi using NicheNet and alternative graph configurations. The table summarizes the predictive performance of CellNavi across different gene graph configurations, including NicheNet and alternative designs, evaluated on the Schmidt dataset. Performance metrics include Top-1 accuracy, Top-5 accuracy, area under the receiver operating characteristic curve (AUROC), area under the precision-recall curve (AUPRC), F1 score, and Matthews Correlation Coefficient (MCC). For sparsified and random graph configurations, three independent graphs were generated and evaluated for each setup.

