## Supplementary Information for "Directing cellular transitions on gene graph-enhanced cell state manifold"

### Supplementary Note 1. Dimensionality considerations for cell state manifold

The choice of optimal dimensionality for representing the cell state manifold is a critical question that warrants further investigation.

In this study, we utilized a 2,048-dimensional coordinate space to represent the manifold of cell states. This dimensionality was chosen based on a combination of theoretical considerations and prior literature, and we presume that the precise dimensionality is not critical for constructing the cell state manifold itself.

The primary desiderata for constructing the manifold are (1) lower dimensionality than the original gene expression space and (2) biological interpretability of distances in the vector space. Lower dimensionality ensures the removal of redundant information, as the intrinsic dimensionality of cellular states is significantly lower than the ~20,000-dimensional gene expression space due to biological constraints and gene co-regulation. Biological interpretability ensures that distances in the manifold correspond to meaningful differences between cell states. For the first criterion, it is sufficient to select a dimensionality smaller than the original space, as the intrinsic dimensionality of the manifold is constrained by inter-gene relationships. For the second criterion, theoretical results such as the Nash embedding theorem (Nash, John (1954). "C1 isometric embeddings." *Annals of Mathematics*. 60(3): 383–396) demonstrate that any manifold of intrinsic dimensionality $n$ can be faithfully embedded in a coordinate space of dimensionality no greater than $2n + 1$. Consequently, choosing a dimensionality larger than the intrinsic manifold dimensionality ensures that geometric properties are preserved. Furthermore, the use of down-sampling-reconstruction pretraining techniques in our model enforces biological meaning in the learned embeddings. Importantly, overestimating the dimensionality does not harm the representation, as redundant dimensions can correlate with the intrinsic ones, allowing for an isometric embedding without distorting biological information.

Previous single-cell foundation models, such as Geneformer, scGPT, and scFoundation, have successfully employed latent spaces of 256 or 512 dimensions to represent cell states across diverse biological systems. However, these models did not focus on recovering individual gene expression profiles from cell embeddings, a task central to our method. We hypothesize that a higher dimensionality is beneficial for achieving such recovery, as it provides greater representational capacity. The dimensionality of 2,048 was chosen as a balance between computational efficiency and the ability to encode complex biological relationships.

Due to resource limitations, we did not conduct a systematic evaluation of different dimensionalities in this study. This remains an exciting avenue for future exploration. Importantly, our framework is flexible and allows the dimensionality to be adjusted based on the dataset and specific application. This adaptability ensures that our approach can accommodate diverse biological scenarios.

### Supplementary Note 2. Choice of down-sampling rate in CellNavi pre-training

The motivation for down-sampling-reconstruction strategy is to address the issue of batch effects caused by varying sequencing depths in single-cell RNA-seq datasets. Differences in sequencing depth often result in the loss of low-expression genes, exacerbating technical variability and causing cells with similar biological identities to cluster separately[1,2]. To mitigate this, we simulate depth variation by selectively masking low-expression genes at controlled down-sampling rates and train the model to reconstruct the original gene expression profiles. We expect this strategy enables the model to learn embeddings that are robust to data sparsity and align cell representations across datasets with differing depths. Below, we elaborate on the rationale behind the selection of the current down-sampling rate and present results exploring the sensitivity of the model to different down-sampling rates.

The down-sampling rate (`ds`) determines the fraction of UMI counts retained, with higher rates leading to more aggressive sparsity. Given the computational resources and time required for pre-training, we empirically selected the down-sampling rate based on an analysis of real-world single-cell RNA-seq datasets.

First, we analyzed the distribution of median UMI counts per cell across 1,487 human single-cell RNA-seq datasets collected from the Human Cell Atlas. As shown in the histogram and empirical cumulative distribution (ECDF) in **Supplementary Fig. 13**, the majority (80%) of datasets exhibit a median UMI count between 1,000 and 10,000, while half the datasets have a median UMI below 3,184. A down-sampling rate of 20 simulates a scenario where datasets with median UMI counts near 10,000 are reduced to approximately 500 UMIs per cell, a level at which meaningful biological interpretation becomes increasingly challenging [3,4]. This analysis suggests a maximum down-sampling rate of 20, a realistic sparsity level that approaches the threshold where data may become biologically uninformative in certain scenarios.

To further examine the impact of down-sampling rates on cell-type resolution, we used a human pancreas scRNA-seq dataset as a representative example (**Supplementary Fig. 14**)[5]. We tested various rates (`ds=5, 10, 20, 30, 50, 100`) and found that dominant cell populations remained distinguishable even at high down-sampling rates, likely due to high-expression marker genes. However, moderate cell types, such as quiescent and activated stellate cells and endothelial cells, began to mix at rates beyond `ds=20`, as shown in **Supplementary Fig. 14**. Based on these observations, we constrained the maximum down-sampling rate to 20 in our experiments to preserve the resolution of rare populations while simulating realistic sparsity levels.

For the final pretraining procedure, we employed a mixed down-sampling strategy with rates ranging from `ds=1` (no down-sampling) to `ds=20` (ds1-20). This approach simulates a continuous variation in sequencing depth across datasets, enhancing the model's ability to generalize while mitigating out-of-distribution challenges between the input (down-sampled data) and the output (original expression profiles). This strategy aligns with principles from robust learning frameworks[6]. Including `ds=1` introduces inherent noise during pretraining, which may act as a form of regularization, further improving model robustness and performance (not tested).

### Supplementary Note 3. Evaluate Performance of Linear Methods in Driver Gene Prediction

Several studies have proposed that linear models may outperform complex deep learning-based methods in cell modeling [7–10]. Here, we evaluated predictive capabilities of different linear models in identifying driver genes under various conditions and compared their performance to that of CellNavi. These baselines included the additive model [7], matrix optimization [7], and logistic regression (LR) [10]. All experiments were conducted using the Norman dataset [11], with a train-test split designed to assess both in-domain (cell states seen during training) and out-of-domain (OOD, cell states unseen during training) generalization. For all methods, the objective was to predict driver genes given transcriptomes from a pair of source and target cells.

#### Methods

The additive model was applied to a double perturbation scenario, where single-perturbations from all but one cluster, stratified using Leiden algorithm, were used for training (the same as in our manuscript) and double-perturbations were used for testing. The double perturbation outcome was simulated as $y_a + y_b - 2c$, where $y_a$ and $y_b$ represent the transcriptomic outcomes of single gene perturbations $g_a$ and $g_b$, respectively, and $c$ is the control. We randomly sampled 100 combinations of $y_a$, $y_b$, and $c$ and computed their averages to construct a $G \times P_{pair}$ matrix, where each row represents a gene, each column represents a perturbation pair, and each cell corresponds to the expression value of the gene upon perturbation of the pair. Testing involved calculating $y_{test} - c_{test}$, then computing the cosine similarity between this difference and each column in the $G \times P_{pair}$ matrix, ranking the perturbation pairs by similarity.

Next, we applied matrix optimization approach to single gene perturbations prediction, we followed a similar methodology to the one described in the reference study [7]. Since our goal is to develop a method that applies to unseen cell states, rather than unseen genes, this approach necessitates some adjustments. Gene expression changes were calculated from the training set, and pseudobulking was performed to create a $\Delta G \times P$ matrix. Testing involved calculating the cosine similarity between $y_{test} - c_{test}$ and the columns of $\Delta G \times P$, ranking the perturbation pairs accordingly.

To explore learnable linear methods, we implemented an L1-regularized logistic regression (LR) model, following previous description [10]. For each pair of control and perturbed cells, we calculated the change in gene expression, concatenated these changes into a vector of length $G$, and used it as input for a logistic regression classifier. The task was framed as multi-class classification, where the model learned to prioritize driver genes through cross-entropy loss. The LR baseline was evaluated under two settings: (1) out-of-domain (OOD), where single-perturbations from unseen cell states were used for testing, and (2) random splitting, where the test set was randomly selected from the same distribution as the training data.

**Results**

For double perturbation prediction, the additive model performed poorly in both in-domain and out-of-domain settings, as summarized in **Supplementary Table 5**. In the in-domain setting, the model achieved only a Top-2 accuracy of 0.0288 and a F1 score of 0.0893. In the out-of-domain setting, the additive model's performance further deteriorated, with a Top-2 accuracy of 0.0000 and a F1 score of 0.0129, significantly underperforming compared to CellNavi across all metrics. These results indicate that the additive model is not a viable alternative for the driver gene prediction task.

For single perturbation prediction (**Supplementary Table 6**), in the OOD setting, the matrix optimization baseline demonstrated poor performance, achieving only 0.0019 Top-1 accuracy. The logistic regression model, while showing improvements over matrix optimization, still underperformed compared to CellNavi. These results suggest that linear models struggle to generalize to unseen cell states, especially for complex, cell-specific perturbation responses. Even with random splitting, the LR model's performance was substantially lower than that of CellNavi in the OOD setting.

The results demonstrate that both model-free and linear methods fail to deliver robust performance in the driver gene prediction task, particularly in the context of cross-cell-state generalization. The additive model and matrix optimization approaches, which rely on predefined rules or similarity measures, lack the capacity to capture the complex, non-linear perturbation effects specific to individual cell states. Even the logistic regression model, while more effective than the matrix optimization baseline, is fundamentally limited in its ability to generalize to unseen cell states due to its linear nature. These findings underscore the need for more sophisticated approaches, such as CellNavi, which leverage non-linear embeddings and cell-specific perturbation effects to achieve significantly higher performance.

### Supplementary Note 4. Extended training information for CellNavi

#### Optional Training Schema for CellNavi to Mitigate Batch Effect

In some cases, cellular transition data for training and testing the DGP in CellNavi may originate from different sequencing platforms or experimental techniques. Such technical discrepancies can introduce batch effects that obscure biological insights. We hence propose two additional finetuning strategies for the option to account for the batch effect.

##### *Cell Coordinate Distribution Alignment*

Batch effects may introduce shifts of the cell coordinate distributions over two batches, due to biologically irrelevant factors which we seek to eliminate. To correct for this bias, we adopt the adversarial generative training strategy[6]. The intuition is to ask the CMM to produce cell coordinates that confuse a batch label classifier, i.e., worsen the batch label classification accuracy. In this way, the cell coordinates are forced to discard any information about the batch it comes from, hence eliminating the distribution shift between the cell coordinates of the two batches.

Specifically, we use a two-layer MLP for the auxiliary batch label classifier model, also called a discriminator. The input of the MLP is the concatenation of the cell coordinates of a pair of cells from the same batch, and the output is a scalar representing the predicted logit for the binary classification of which batch does the pair come from. The loss function for the discriminator is hence the binary cross-entropy loss:

$$\mathcal{L}_{\mathrm{discr}} = \mathrm{CE}\left(\mathrm{DISCR}\left(\mathrm{CONCAT}\left(\mathbf{CRD}(\mathbf{X}_{\mathrm{src}}), \mathbf{CRD}(\mathbf{X}_{\mathrm{tgt}})\right)\right), y_{\mathrm{batch}}\right),$$

where $y_{\mathrm{batch}}$ represents the binary label indicating whether the input cell pair $\mathbf{X}_{\mathrm{src}}$ and $\mathbf{X}_{\mathrm{tgt}}$ comes from the test batch (in contrast to the training batch). The discriminator is asked to minimize the loss in order to try to extract any batch-related information from the cell coordinates, while the CMM is asked to maximize the loss with the help of the discriminator in order to reduce batch-related information to eliminate the batch effect. In practice, the discriminator is optimized alternately with the optimization of CMM.

***Finetuning Stabilization by Cell Label Classifier***

The adversarial training process is known unstable, and could degenerate the cell coordinates to a trivial representation that does not contain even biologically relevant information. To combat this, we also introduce an auxiliary cell label classifier, which uses the cell coordinates produced by the CMM to predict a biologically related cell label. To allow accurate prediction, the CMM is enforced to keep biologically related information in the cell coordinates. The cell label classifier is also a two-layer MLP, and is trained by the following loss function:

$$\mathcal{L}_{\text{cell}} = \text{CE}\big(\text{CELLCLS}(\mathbf{CRD}(\mathbf{X})), y_{\text{cell\_label}}\big),$$

where $y_{\text{cell\_label}}$ denotes the cell label of cell $\mathbf{X}$. The loss is applied exclusively on the test data set, and optimizes both the CMM and the cell label classifier $\text{CELLCLS}$.

***Combined Loss for Finetuning***

The overall loss function for the driver gene prediction finetuning in the case of batch effect is:

$$\mathcal{L}'_{\text{driver\_gene}} = \alpha_{\text{driver\_gene}}\mathcal{L}_{\text{driver\_gene}} + \alpha_{\text{cell}}\mathcal{L}_{\text{cell}} - \alpha_{\text{discr}}\mathcal{L}_{\text{discr}},$$

where $\alpha_{\text{driver\_gene}}$ , $\alpha_{\text{cell}}$ and $\alpha_{\text{discr}}$ are hyperparameters that control the weights of the respective losses. This loss optimizes the CMM, DGP, and the cell label classifier jointly, in alternation with the optimization of the discriminator which minimizes $\mathcal{L}_{\text{discr}}$, as mentioned. Note that the CMM aims to maximize $\mathcal{L}_{\text{discr}}$ (hence the minus sign) with the help of the discriminator. This multi-objective setup promotes alignment across batches while maintaining robust biological representation in the data.

**Training details**

The pre-training of CellNavi was conducted with 32 A100. Fine-tuning and inference were conducted using single NVIDIA A100 GPU (80 GB PCIe) and single Intel Xeon Gold 6342 CPU (24 cores, 2.80 GHz). The fine-tuning phase, performed on task-specific datasets ranging from 28,453 to 111,445 cells, required 14 to 40 hours depending on the dataset size, with a batch size of 128. For inference, the model processes each cell with a speed of approximately 0.38 seconds.

**Supplementary Table 1. Comparison of Method Performance for Driver Gene Prediction on the Schmidt and Norman Datasets**

| | Top-1 accuracy | Top-5 accuracy | AUROC (macro) | AUPRC (macro) | F1 score (macro) | MCC |
|---|---|---|---|---|---|---|
| **Schmidt dataset** | | | | | | |
| CellNavi | **0.6206** | **0.7326** | **0.8745** | **0.4991** | **0.6154** | **0.6155** |
| GEARS | 0.0520 | 0.2110 | 0.5170 | 0.0307 | 0.0276 | 0.0317 |
| SCENIC | 0.0930 | 0.3482 | 0.6274 | 0.0689 | 0.0493 | 0.0667 |
| DGE | 0.0693 | 0.2104 | 0.6295 | 0.0790 | 0.0483 | 0.0578 |
| GENIE3 | 0.1389 | 0.4941 | 0.5785 | 0.1000 | 0.0084 | 0.0842 |
| GRNBoost2 | 0.1049 | 0.2718 | 0.6106 | 0.0788 | 0.0787 | 0.0843 |
| RENGE | 0.0358 | 0.2738 | 0.5133 | 0.0468 | 0.0034 | 0.0126 |
| **Norman dataset (single perturbation)** | | | | | | |
| CellNavi | **0.4900** | **0.6330** | **0.8884** | **0.4540** | **0.4219** | **0.4816** |
| GEARS | 0.1140 | 0.2970 | 0.5713 | 0.0228 | 0.1005 | 0.0963 |
| SCENIC | 0.0562 | 0.1420 | 0.5535 | 0.0267 | 0.0437 | 0.0439 |
| DGE | 0.0319 | 0.0931 | 0.5474 | 0.0137 | 0.0074 | 0.0098 |
| GENIE3 | 0.0865 | 0.1911 | 0.5909 | 0.0318 | 0.0693 | 0.0739 |
| GRNBoost2 | 0.0313 | 0.0902 | 0.5301 | 0.0183 | 0.0239 | 0.0363 |
| RENGE | 0.0516 | 0.3107 | 0.5836 | 0.0636 | 0.0160 | 0.0062 |

The table summarizes the predictive performance of CellNavi and alternative methods in Schmidt dataset and Norman dataset (single perturbation). Metrics reported include Top-1 and Top-5 accuracy, F1 score, area under the receiver operating characteristic curve (AUROC), area under the precision-recall curve (AUPRC), and Matthews Correlation Coefficient (MCC).

**Supplementary Table 2. Cross-Validation Results for Driver Gene Prediction on the Schmidt Dataset**

| | Top-1 accuracy | Top-5 accuracy | AUROC | AUPRC | F1 score | MCC |
|---|---|---|---|---|---|---|
| *Train: Re-stimulated T cells. Test: Resting T cells (used in Figure 2)* | | | | | | |
| CellNavi | **0.6206** | **0.7326** | **0.8745** | **0.4991** | **0.6154** | **0.6155** |
| SCENIC | 0.0930 | 0.3482 | 0.6274 | 0.0689 | 0.0493 | 0.0667 |
| GEARS | 0.0520 | 0.2110 | 0.5170 | 0.0307 | 0.0276 | 0.0317 |
| DGE | 0.0693 | 0.2104 | 0.6295 | 0.0790 | 0.0483 | 0.0578 |
| *Train: Resting T cells. Test: Re-stimulated T cells* | | | | | | |
| CellNavi | **0.4020** | **0.5313** | **0.8021** | **0.2809** | **0.4223** | **0.4013** |
| SCENIC | 0.0672 | 0.2366 | 0.5936 | 0.0509 | 0.0599 | 0.0492 |
| GEARS | 0.1021 | 0.2747 | 0.5388 | 0.0362 | 0.0554 | 0.0739 |
| DGE | 0.0722 | 0.2021 | 0.6237 | 0.0345 | 0.0343 | 0.0598 |

The table presents cross-validation results comparing CellNavi with alternative on the Schmidt dataset. Performance metrics include Top-1 accuracy, Top-5 accuracy, area under the receiver operating characteristic curve (AUROC), area under the precision-recall curve (AUPRC), F1 score, and Matthews Correlation Coefficient (MCC). Two experimental conditions are shown: (1) training on re-stimulated T cells and testing on resting T cells, and (2) training on resting T cells and testing on re-stimulated T cells.

**Supplementary Table 3. Cross-Validation Results for Driver Gene Prediction on the Norman Dataset**

| | Top-1 accuracy | Top-5 accuracy | AUROC | AUPRC | F1 score | MCC |
|---|---|---|---|---|---|---|
| *Split 1 (used in Figure 2)* | | | | | | |
| CellNavi | **0.4900** | **0.6330** | **0.8884** | **0.4540** | **0.4219** | **0.4816** |
| SCENIC | 0.1140 | 0.2970 | 0.5713 | 0.0228 | 0.1005 | 0.0963 |
| GEARS | 0.0562 | 0.1420 | 0.5535 | 0.0267 | 0.0437 | 0.0439 |
| DGE | 0.0319 | 0.0931 | 0.5474 | 0.0137 | 0.0103 | 0.0098 |
| *Split 2* | | | | | | |
| CellNavi | **0.8632** | **0.9259** | **0.9723** | **0.8496** | **0.1384** | **0.7074** |
| SCENIC | 0.0019 | 0.0061 | 0.4304 | 0.0108 | 0.0005 | 0.0063 |
| GEARS | 0.0226 | 0.0600 | 0.5416 | 0.0203 | 0.0331 | 0.0147 |
| DGE | 0.7691 | 0.7769 | 0.9164 | 0.6053 | 0.8002 | 0.5616 |
| *Split 3* | | | | | | |
| CellNavi | **0.4902** | **0.6997** | **0.8960** | **0.4438** | **0.3821** | **0.4834** |
| SCENIC | 0.0529 | 0.1252 | 0.5456 | 0.0249 | 0.0409 | 0.0509 |
| GEARS | 0.1647 | 0.3425 | 0.5335 | 0.0183 | 0.1262 | 0.1500 |
| DGE | 0.0412 | 0.1309 | 0.5969 | 0.0205 | 0.0129 | 0.0341 |
| *Split 4* | | | | | | |
| CellNavi | **0.4609** | **0.6405** | **0.8676** | **0.3668** | **0.4224** | **0.4547** |
| SCENIC | 0.0221 | 0.0831 | 0.5265 | 0.0182 | 0.0175 | 0.0336 |
| GEARS | 0.1429 | 0.3007 | 0.5582 | 0.0221 | 0.1332 | 0.1259 |
| DGE | 0.0238 | 0.0699 | 0.5427 | 0.0204 | 0.0223 | 0.0167 |
| *Split 5* | | | | | | |
| CellNavi | **0.5944** | **0.7857** | **0.9118** | **0.5336** | **0.1671** | **0.5581** |
| SCENIC | 0.0734 | 0.1335 | 0.7148 | 0.0341 | 0.0478 | 0.0786 |
| GEARS | 0.2019 | 0.3270 | 0.5429 | 0.0202 | 0.1562 | 0.1605 |
| DGE | 0.2564 | 0.4126 | 0.8245 | 0.1345 | 0.1632 | 0.1981 |

The table presents cross-validation results comparing CellNavi with alternative on the Norman dataset, using single perturbation experiments. Performance metrics include Top-1 accuracy, Top-5 accuracy, area under the receiver operating characteristic curve (AUROC), area under the precision-recall curve (AUPRC), F1 score, and Matthews Correlation Coefficient (MCC). Split 1 is the same split as used in Figure 2.

**Supplementary Table 4: Driver genes related to cytokine secretion and Th2 differentiation**

| Gene | Function | Reference |
|---|---|---|
| CD28 | IL2 secretion, IFNG secretion | doi: 10.1126/science.abj4008<br>doi: 10.4049/jimmunol.1500707<br>https://www.nature.com/articles/ni0101_37 |
| VAV1 | IL2 secretion, IFNG secretion | doi: 10.1126/science.abj4008<br>https://www.nature.com/articles/374474a0<br>https://doi.org/10.4049/jimmunol.177.8.5024 |
| CD27 | IFNG secretion | doi: 10.1126/science.abj4008<br>https://doi.org/10.1016/j.immuni.2024.01.011 |
| IL9R | IFNG secretion | doi: 10.1126/science.abj4008<br>https://www.nature.com/articles/s41586-022-04801-2 |
| GATA3 | Th2 differentiation | doi: 10.1126/science.abj4008<br>https://doi.org/10.1016/j.cell.2018.11.044<br>https://www.nature.com/articles/ncomms2260 |

### Supplementary Table 5. Comparison of predictive performance between the linear model and CellNavi for double perturbation prediction

| | Top-2 accuracy | Top-5 accuracy | Top-10 accuracy | F1 score | AUROC | AUPRC |
|---|---|---|---|---|---|---|
| **In domain (cell state seen)** | | | | | | |
| Additive model | 0.0288 | 0.1550 | 0.1969 | 0.0893 | 0.5063 | 0.0090 |
| CellNavi | 0.2094 | 0.3627 | 0.4881 | 0.4100 | 0.8100 | 0.3938 |
| **Out of domain (cell state unseen)** | | | | | | |
| Additive model | 0.0000 | 0.0039 | 0.0039 | 0.0129 | 0.5412 | 0.0098 |
| CellNavi | 0.1358 | 0.2825 | 0.3760 | 0.3520 | 0.7916 | 0.3078 |

The table summarizes the predictive performance of the additive model and CellNavi in two scenarios: **(1) in-domain**, where the model is tested on double perturbations of cell states seen during training, and **(2) out-of-domain**, where the model is tested on double perturbations of cell states unseen during training. Metrics reported include Top-2, Top-5, and Top-10 accuracy, F1 score, area under the receiver operating characteristic curve (AUROC), and area under the precision-recall curve (AUPRC).

### Supplementary Table 6. Comparison of predictive performance between the linear models and CellNavi for single perturbation prediction

| | Top-1 accuracy | F1 score | AUROC | AUPRC |
|---|---|---|---|---|
| **Out of domain (cell state unseen)** | | | | |
| Matrix optimization | 0.0019 | 0.2326 | 0.5230 | 0.0114 |
| Linear regression | 0.2078 | 0.1711 | 0.7169 | 0.1736 |
| CellNavi | 0.4900 | 0.4219 | 0.8884 | 0.4540 |
| **Random split** | | | | |
| Linear regression | 0. 3085 | 0.2592 | 0.8024 | 0.2604 |

**Out-of-domain**: The model is tested on single perturbations from cell states unseen during training. **Random split**: The test set is randomly selected from the perturbation dataset. Metrics reported include Top-1 accuracy, F1 score, area under the receiver operating characteristic curve (AUROC), and area under the precision-recall curve (AUPRC).

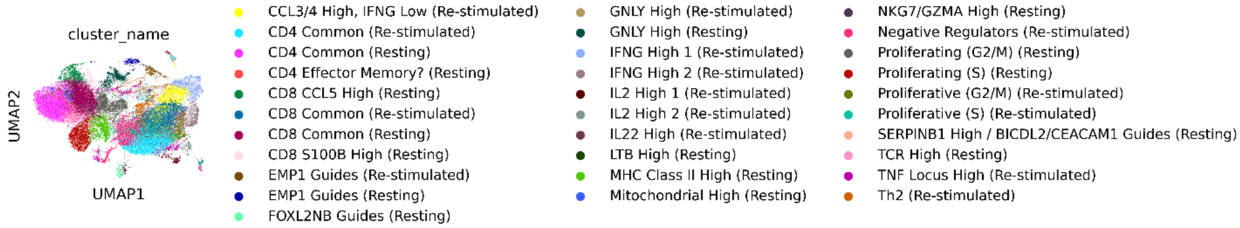


**Supplementary Fig. 1** UMAP visualization of perturbed resting T cells and restimulated T cells sequenced by Schmidt et al, colored by cell clusters identified in the original study.

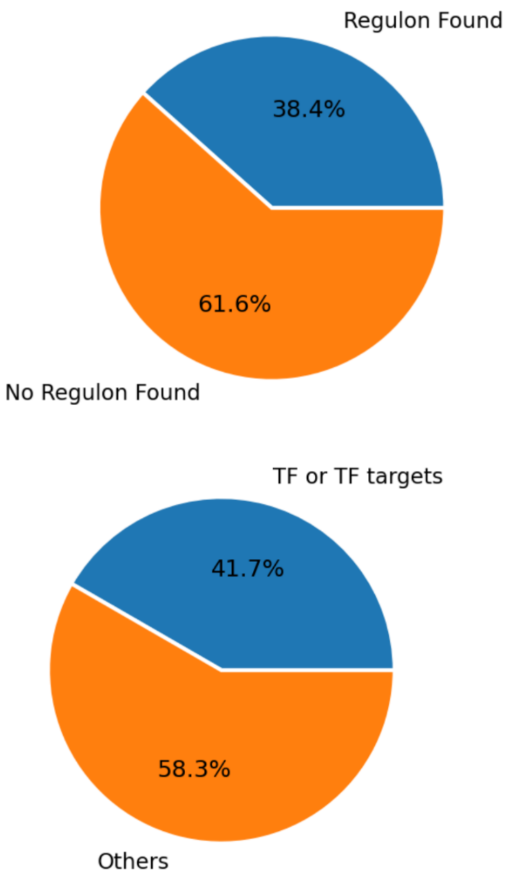


**Supplementary Fig. 2** Compatibility of the test dataset to SCENIC/SCENIC+. Upper panel: 61.6% of samples cannot be predicted by SCENIC/SCENIC+ due to missing regulons. Bottom panel: 58.3% of candidate driver genes are excluded because they are neither transcription factor (TF) nor TF-target genes.

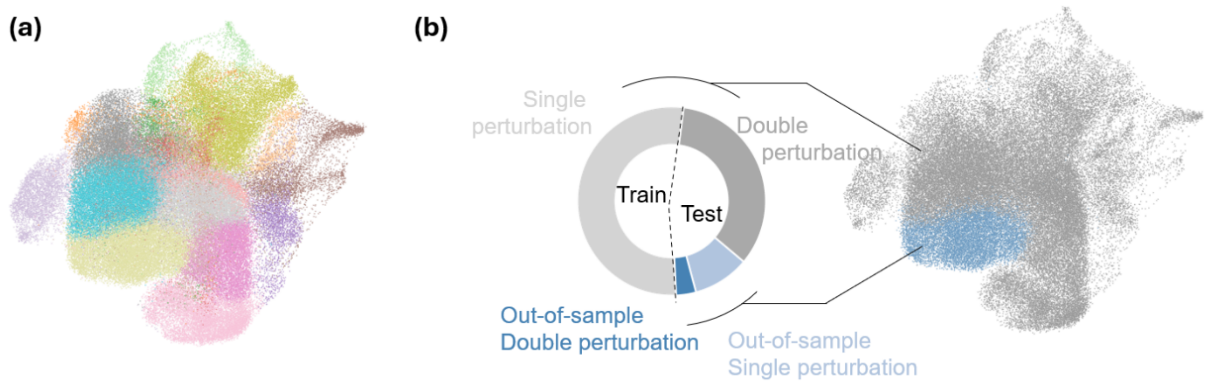


**Supplementary Fig. 3 UMAP visualization of the train-test data split for the Norman dataset. (a)** Cells were stratified into distinct states using the unsupervised Leiden algorithm, with each state represented by a different color. **(b)** A specific cell cluster was selected as the test set, while the remaining clusters were used for training. To ensure rigorous evaluation, all multi-gene perturbations were excluded from training. Consequently, the training set (light gray) consisted only of single-gene perturbations within certain clusters, while the test set was divided into: 1) single-gene perturbations from the held-out cluster (light blue), 2) double-gene perturbations from the held-out cluster (dark blue), and 3) double-gene perturbations from the training clusters (dark gray). Results reported in the main text are derived from the held-out cluster (blue cluster in the UMAP). To further ensure fairness, the test cluster was shuffled (similar to cross-validation) to obtain robust and unbiased results (**Supplementary Table 3**).

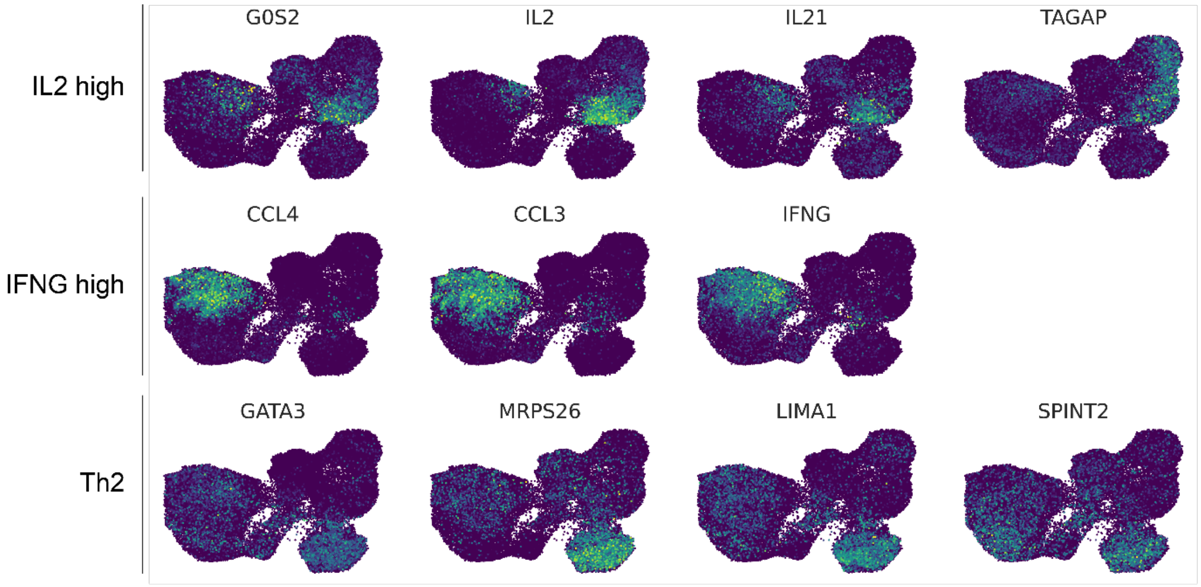


**Supplementary Fig. 4 Transcriptional changes of canonical marker genes in IL2-high, IFNG-high, and Th2 cell types.** IL2-high marker genes**:** *G0S2, IL2, IL21, and TAGAP*. IFNG-high marker genes: *IFNG, CCL3, CCL4, and CCL3L3*. Th2 marker genes: *GATA3, MRPS26, LIMA1 and SPINT2*.

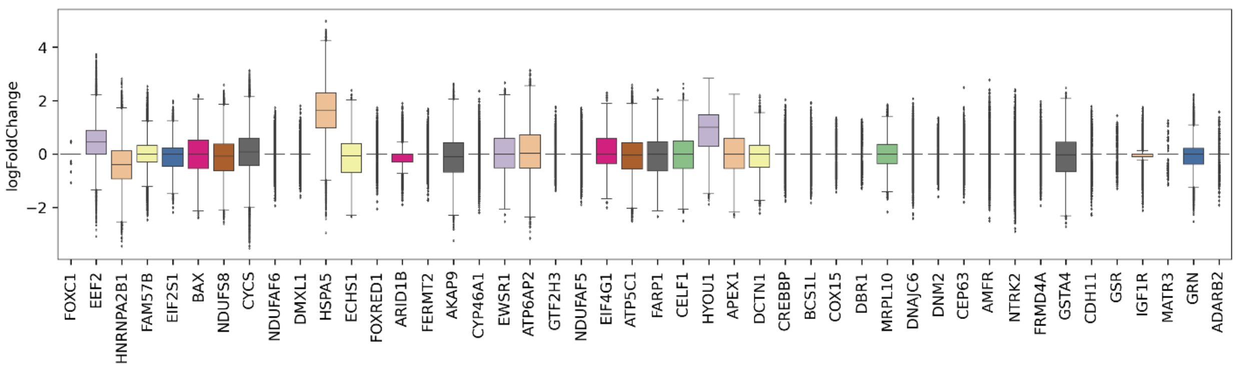


**Supplementary Fig. 5** Expression changes for the top-20 predicted genes across cell pairs. Center line, median; box limits, upper and lower quartiles; whiskers, 1.5x interquartile range; points, outliers. *n* = 47,437.

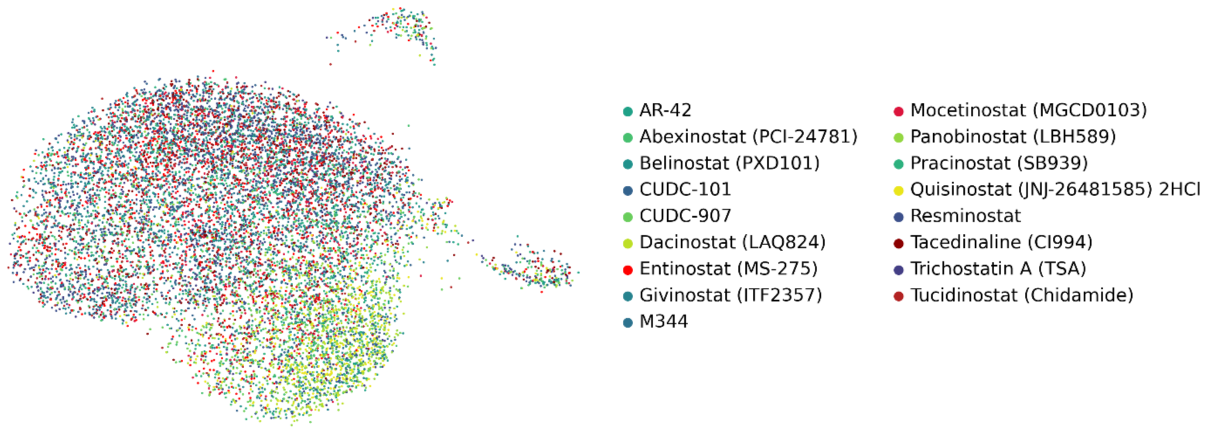


**Supplementary Fig. 6** UMAP visualization of single-cell transcriptomic profiles from K562 cells treated with 17 distinct HDAC inhibitors.

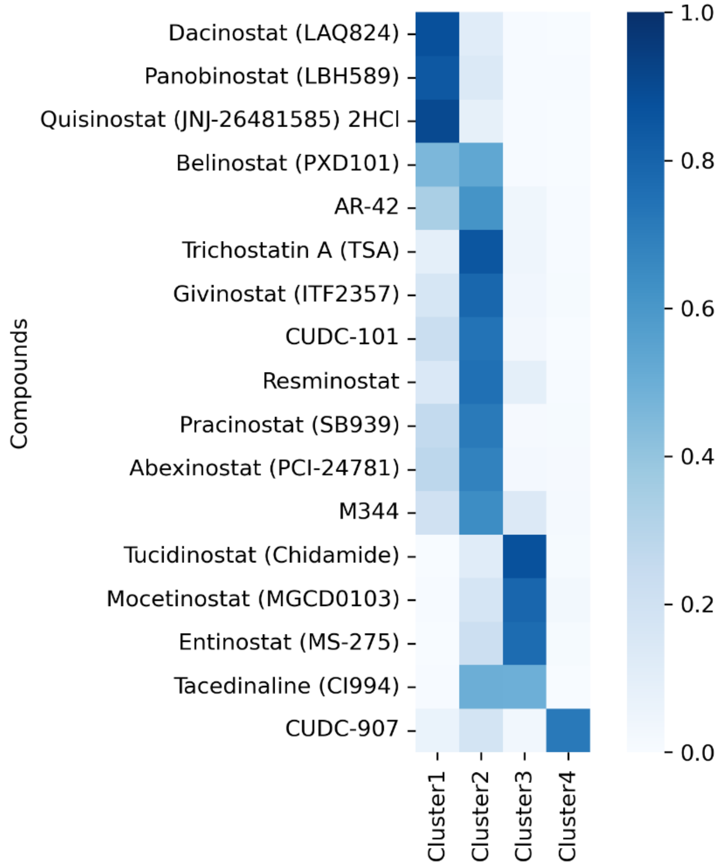


**Supplementary Fig. 7** The percentage of cells treated with different drug compounds within each cluster identified in Fig. 5a

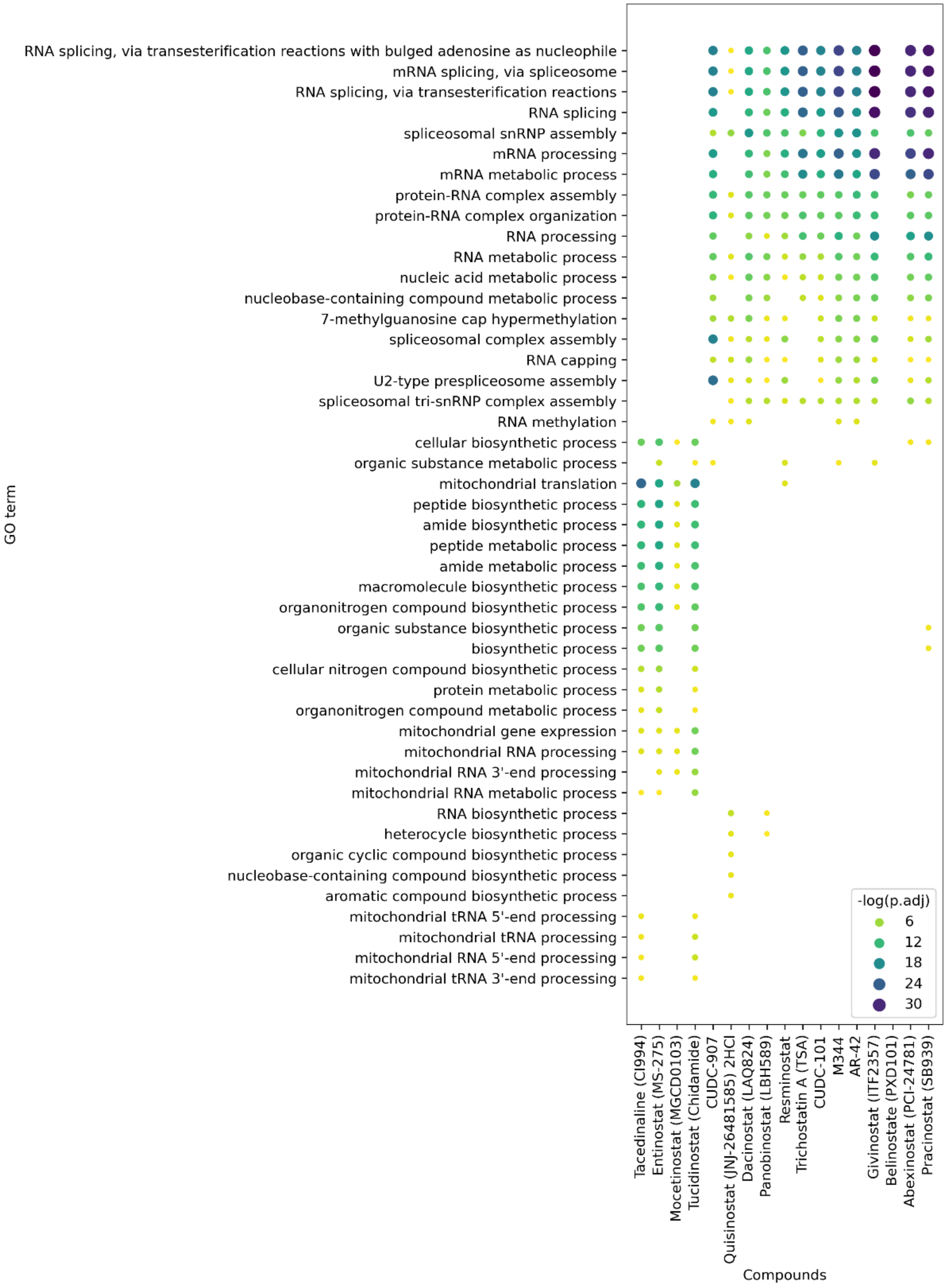


**Supplementary Fig. 8** Gene Ontology (GO) enrichment analysis for each treatment group. The size and darkness of the dots correlate negatively with the adjusted *p*-value (one-sided Fisher's exact test with Benjamini–Hochberg correction for multiple comparisons).

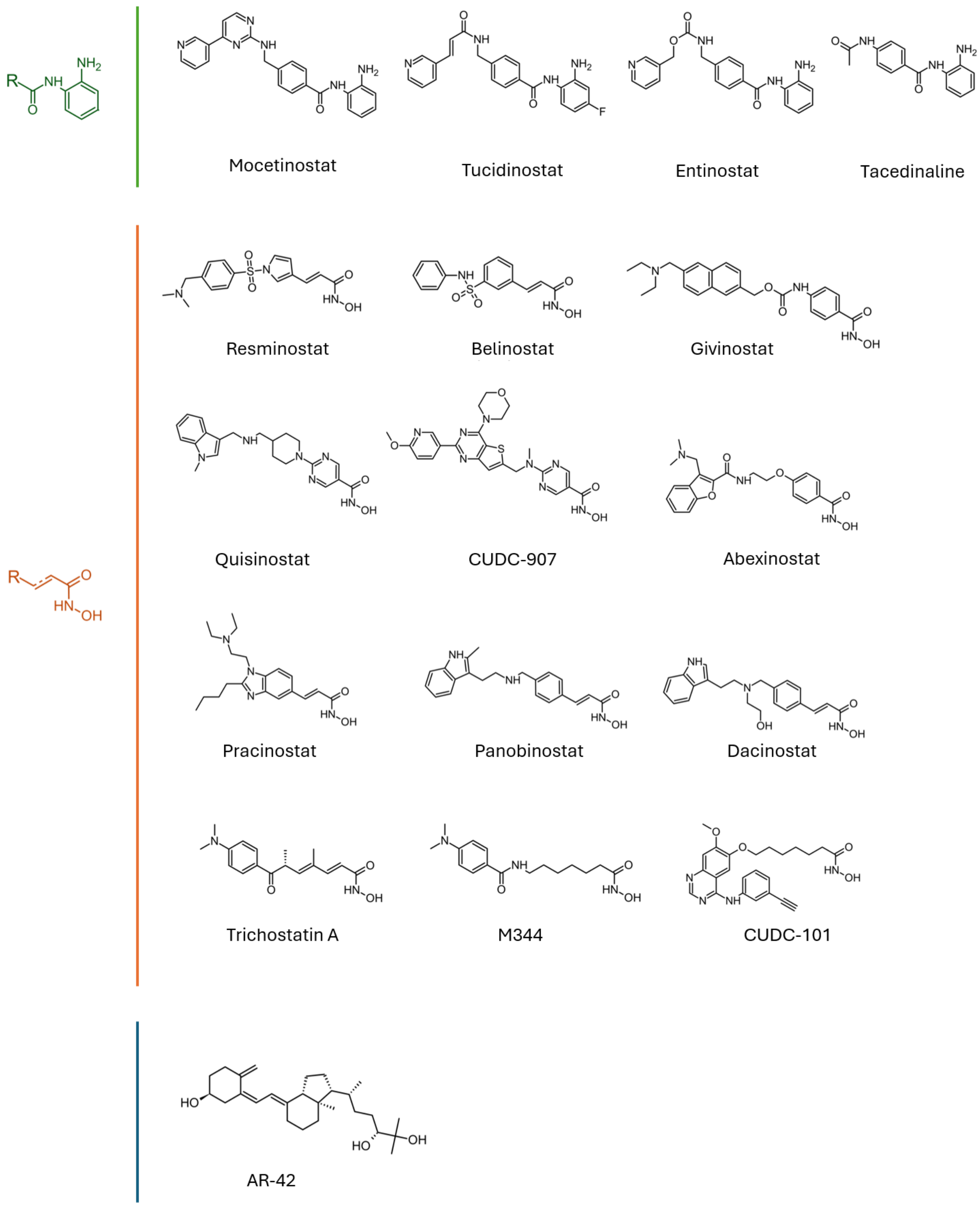


**Supplementary Fig. 9** Molecular structures of the 17 HDAC inhibitors. Except for AR-42, all other molecules share two distinct types of warheads.

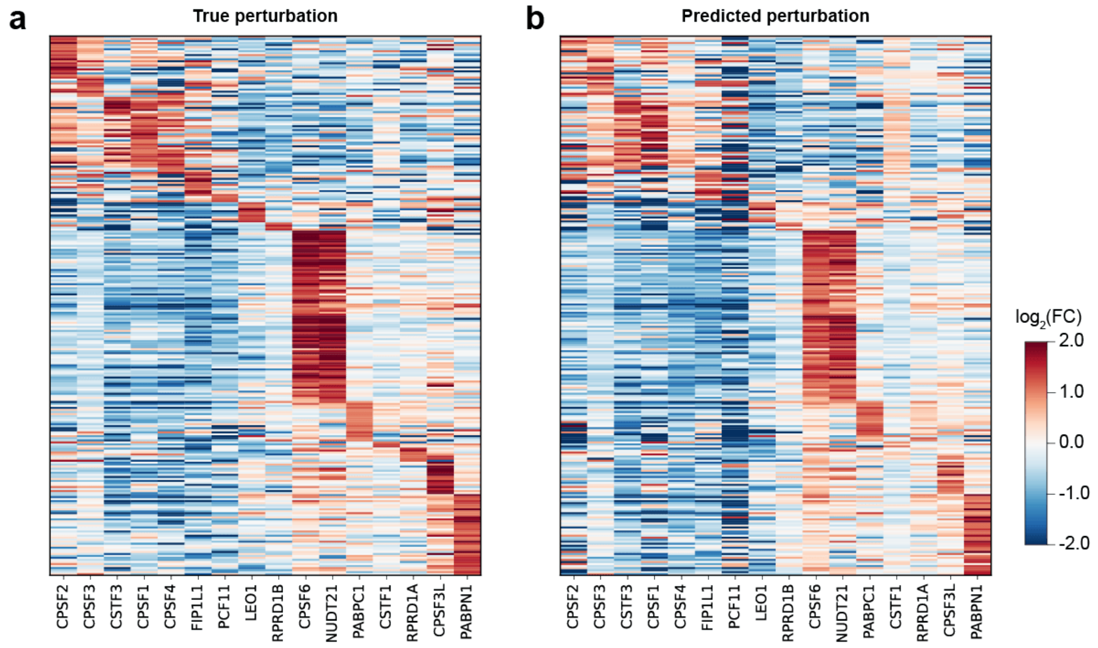


**Supplementary Fig. 10** Heatmap showing gene expression across different perturbation groups for K562 cells. Rows represent genes, and columns represent true perturbations **(a)** or predicted perturbations by CellNavi **(b)**.

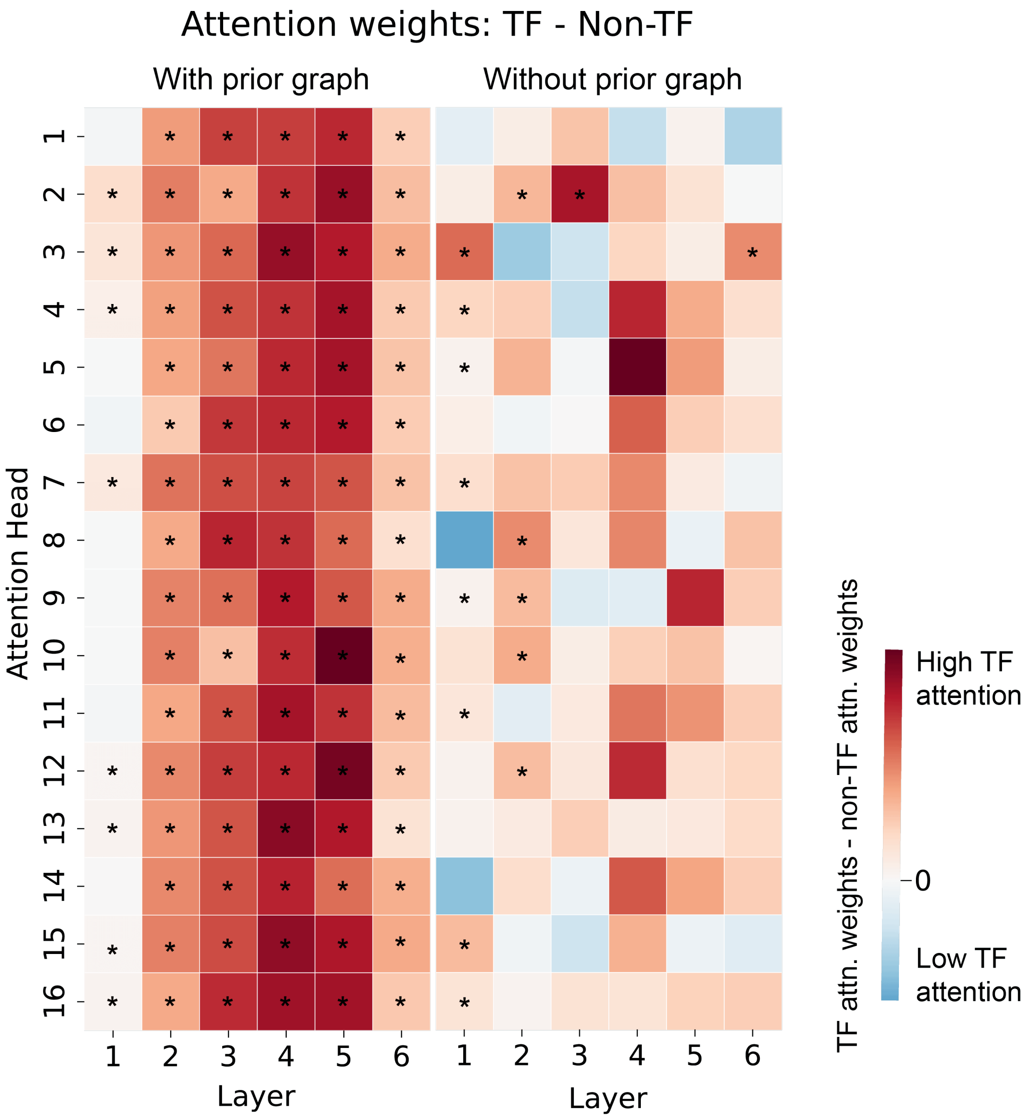


**Supplementary Fig. 11. Differential attention weights between TFs and non-TFs.** The heatmap displays the contrast in attention weights directed towards transcription factors (TFs) versus non-transcription factors (non-TFs) when the prior gene-gene interaction graph is included (left) or excluded (right). The color gradient represents the magnitude of the difference in attention weights. In the presence of the gene-gene graph, 88 attention heads demonstrate a statistically significant bias towards TFs, while without the graph, 15 attention heads show this preference ($P < 0.05$, two-sided Wilcoxon rank sum test, with false discovery rate (FDR) adjustment using the Benjamini–Hochberg procedure). $n$ = 4,067.

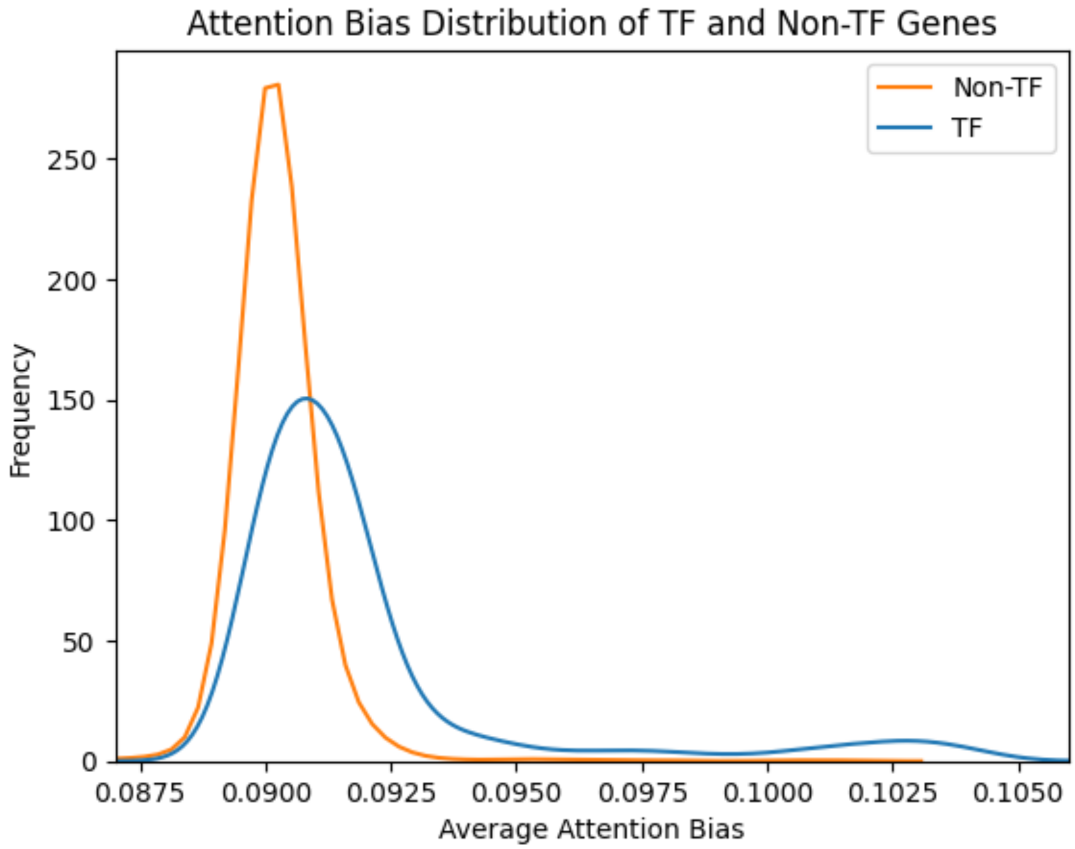


**Supplementary Fig. 12.** The attention bias learned from the gene graph exhibited a notably higher magnitude for TFs compared to non-TFs in distribution ($P < 0.05$, two-sided Wilcoxon rank sum test). To aid visualization, the sample sizes for both TFs and non-TFs have been equalized through random sampling.

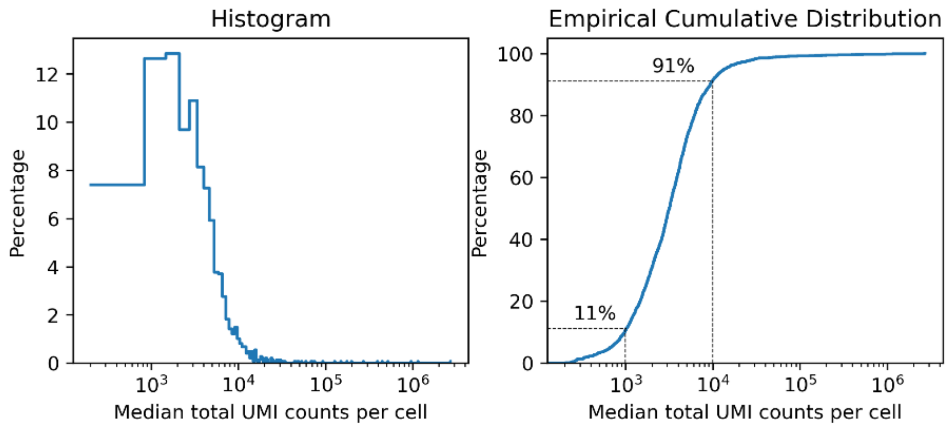


**Supplementary Fig. 13. Distribution of median UMI counts per cell across 1,487 human single-cell RNA-seq datasets.** Left: the histogram of median total UMI counts, highlighting the variability in sequencing depth across datasets. Right: the empirical cumulative distribution (ECDF) of median UMI counts. 91% of datasets have a median UMI count below 10,000, while 11% fall below 1,000.

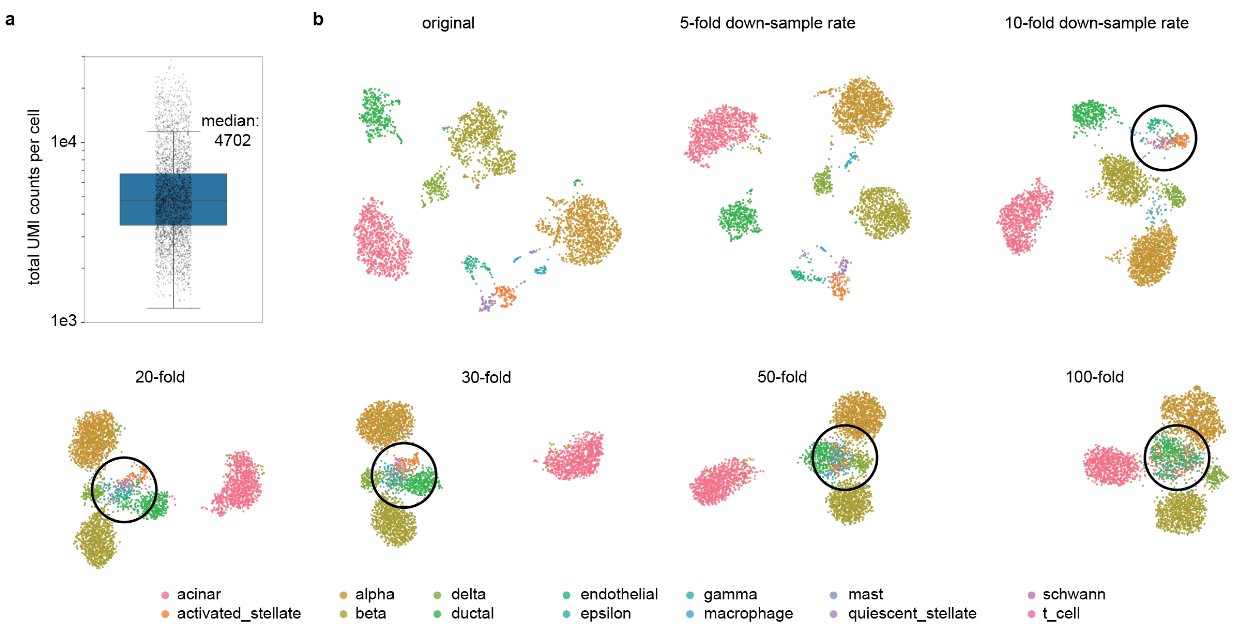


**Supplementary Fig. 14. UMI counts distribution and cell-type clustering at varying down-sampling rates.** (a) UMI counts distribution. Center line, median; box limits, upper and lower quartiles; whiskers, 1.5x interquartile range. *n* = 3,605 (GSE84133 human3). (b) UMAP visualization of the dataset at varying down-sampling rates. Cells are colored by original cell-type annotations. Black circles highlight cell types that start to mix at higher down-sampling rates.
